# Second Heart Field-derived Cells Contribute to Angiotensin II-mediated Ascending Aortopathies

**DOI:** 10.1101/2020.02.02.930917

**Authors:** Hisashi Sawada, Yuriko Katsumata, Hideyuki Higashi, Chen Zhang, Yanming Li, Stephanie Morgan, Lang H. Lee, Sasha A. Singh, Jeff Z. Chen, Michael K. Franklin, Jessica J. Moorleghen, Deborah A. Howatt, Debra L. Rateri, Ying H. Shen, Scott A. LeMaire, Masanori Aikawa, Mark W. Majesky, Hong S. Lu, Alan Daugherty

## Abstract

**Background:** The ascending aorta is a common location for aneurysm and dissection. This aortic region is populated by a mosaic of medial and adventitial cells that are embryonically derived from either the second heart field (SHF) or the cardiac neural crest. SHF-derived cells populate areas that coincide with the spatial specificity of thoracic aortopathies. The purpose of this study was to determine whether and how SHF-derived cells contribute to ascending aortopathies.

**Methods:** Ascending aortic pathologies were examined in patients with sporadic thoracic aortopathies and angiotensin II (AngII)-infused mice. Ascending aortas without overt pathology from AngII-infused mice were subjected to mass spectrometry assisted proteomics, and molecular features of SHF-derived cells were determined by single cell transcriptomic analyses. Genetic deletion of either low-density lipoprotein receptor-related protein 1 (*Lrp1*) or transforming growth factor-β receptor 2 (*Tgfbr2*) in SHF-derived cells was conducted to examine the impact of SHF-derived cells on vascular integrity.

**Results:** Pathologies in human ascending aortic aneurysmal tissues were predominant in outer medial layers and adventitia. This gradient was mimicked in mouse aortas following AngII infusion that was coincident with the distribution of SHF-derived cells. Proteomics indicated that brief AngII infusion, prior to overt pathology, evoked downregulation of SMC proteins and differential expression of extracellular matrix proteins, including several LRP1 ligands. LRP1 deletion in SHF-derived cells augmented AngII-induced ascending aortic aneurysm and rupture. Single cell transcriptomic analysis revealed that brief AngII infusion decreased *Lrp1* and *Tgfbr2* mRNA abundance in SHF-derived cells and induced a unique fibroblast population with low abundance of *Tgfbr2* mRNA. SHF-specific *Tgfbr2* deletion led to embryonic lethality at E12.5 with dilatation of the outflow tract and retroperitoneal hemorrhage. Integration of proteomic and single cell transcriptomics results identified plasminogen activator inhibitor 1 (PAI1) as the most increased protein in SHF-derived SMCs and fibroblasts during AngII infusion. Immunostaining revealed a transmural gradient of PAI1 in both ascending aortas of AngII-infused mice and human ascending aneurysmal aortas that mimicked the gradient of medial and adventitial pathologies.

**Conclusion:** SHF-derived cells exert a critical role in maintaining vascular integrity through LRP1 and TGF-β signaling associated with increases of aortic PAI1.

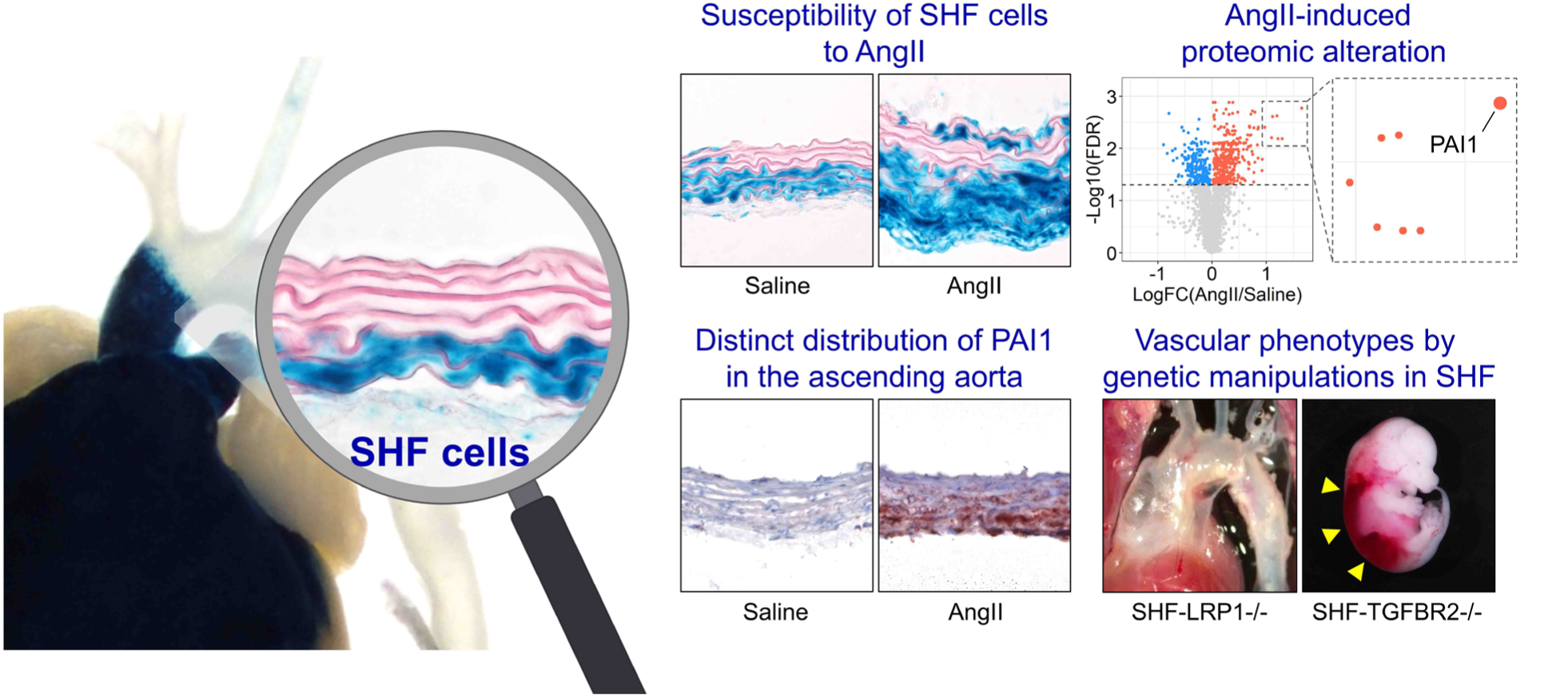

**Clinical Perspective:** *What is new?:*

- SHF-derived SMCs and fibroblasts associate with AngII-induced aortic pathologies.
- AngII induces a distinct fibroblast sub-cluster that is less abundant for mRNAs related to major extracellular components and TGFβ ligands and receptors, but more abundant for proliferative genes.
- TGFBR2 deletion in SHF-derived cells are embryonic lethal with significant dilatation of the outflow tract in mice.
- SHF-specific deletion of LRP1 leads to aortic pathologies in mice, supporting the importance of SHF-derived cells in maintaining ascending aortic wall integrity.

*What are the clinical implications?:*

- Heterogeneity of the embryonic origins of SMCs and fibroblasts contributes to complex mechanisms of vasculopathy formation, which should be considered when investigating the pathogenesis of thoracic aortopathies.

## Introduction

Thoracic aortopathies are a spectrum of lethal diseases associated with aneurysm, dissection, and rupture.^1, 2^ Despite the devastating consequences of these diseases, current medical therapy has limited efficacy. Therefore, there is an urgent need to investigate mechanisms of thoracic aortopathies to facilitate development of effective therapeutics.

Aortopathies occur throughout the thoracic aorta with the ascending aorta being a frequently affected region.^3, 4^ Aortic remodeling, including medial and adventitial thickening, is a histological feature of thoracic aortopathies that shows a transmedial gradient towards the outer medial layers and the adjacent adventitia.^5–10^ Therefore, thoracic aortopathies have regional and transmural specificities with outer medial layers in the ascending aorta being a disease-prone location. However, the mechanism by which outer medial layers of the ascending aorta are prone to aortopathy remains unclear.

The major cellular components of the aortic wall are smooth muscle cells (SMCs) and fibroblasts (FBs). These two cell types are thought to exert a pivotal role in the pathogenesis of thoracic aortopathies.^11^ SMCs in the ascending aorta are derived from two embryonic origins: cardiac neural crest (CNC) and second heart field (SHF).^12^ Select FBs are also derived from the CNC and SHF.^13^ Lineage tracking studies have demonstrated that CNC-and SHF-derived SMCs occupy inner and outer medial layers of the ascending aorta, respectively.^13^ The region-specific distribution of thoracic aortopathies coincides with the distribution of SMCs from the SHF origin. Therefore, we hypothesized that SHF-derived cells play an important role in ascending aortopathies. To test this hypothesis, we investigated whether SHF-derived cells colocalized with the evolution of aortic pathology produced by angiotensin II (AngII) infusion in mice, and investigated the molecular basis of AngII-mediated ascending aortopathies using proteomic and single cell transcriptomic approaches. Our findings using omics analyses were validated by two SHF-specific genetic deletion studies in mice.

## Methods

Additional detailed methods are presented in **Supplemental Methods**. Numerical data are available in **Supplemental Excel File I-III**. All raw data and analytical methods are available from the corresponding author upon appropriate request.

### Human aortic samples

Ascending aortas were acquired from patients undergoing aortic surgery for sporadic thoracic aortic aneurysms at Baylor College of Medicine (n=10, mean age=66±7 years, 5 males and 5 females, mean aortic diameters=5.3±0.4 cm). Aortic tissues were fixed with formalin (10% wt/vol) and then incubated with ethanol (70% vol/vol), as described previously.^14^ Subsequently, tissues were embedded in paraffin blocks, and sliced into 5 µm sections. The protocol for collecting human tissue samples was approved by the Institutional Review Board at Baylor College of Medicine.

### Mice

The following mice were purchased from The Jackson Laboratory (**Supplemental Table I**): ROSA26R*^LacZ^* (#003474), ROSA26R*^mT/mG^* (#007676), *Lrp1* floxed (#012604), *Tgfbr2* floxed (#012603), and Wnt1-*Cre* (#022501, also known as B6-Wnt1-*Cre*2). Mef2c-*Cre* mice (#030262) were purchased from the Mutant Mouse Resource and Research Center. For cell tracking studies of CNC and SHF origins, either Wnt1- or Mef2c-*Cre* male mice were bred to ROSA26R*^LacZ^* female mice, respectively. For the single cell RNA sequencing (scRNAseq), Mef2c-*Cre* male mice were bred to ROSA26R*^mT/mG^* female mice. Mef2c-*Cre* male mice were crossbred to *Lrp1* floxed female mice to delete *Lrp1* in SHF-derived cells. Mef2c-*Cre* male mice were also used for lineage-specific deletion of *Tgfbr2* in SHF-derived cells. Because of a modest induction of AngII-induced thoracic aortic aneurysms (TAAs) in female mice, only male mice were studied.^15^ All experiments were approved by the IACUC at either the University of Kentucky or Baylor College of Medicine in accordance with the guidelines of the National Institutes of Health.

### Subcutaneous infusion using an osmotic pump

After random assignment, either saline or AngII (1,000 ng/kg/min, H-1705, Bachem) was infused via a subcutaneously implanted osmotic pump (Alzet model 2001 for 3 days and model 2004 for 28 days of infusion, respectively, Durect) into male mice at 10 to 14 weeks of age.^16^ Surgical staples, used to close incision sites, were removed 7 days after surgery. Post-operative pain was alleviated by application of topical lidocaine cream (4% wt/wt, LMX4, #0496-0882-15, Eloquest Healthcare, Inc).

### Statistical analyses

Non-omics data are presented as the mean ± standard error of the mean (a diamond-shaped symbol with error bars) or the median and 25^th^/75^th^ percentiles (a box plot). Normality and homogeneity of variance were assessed by Shapiro-Wilk and Brown-Forsythe tests, respectively. Because the original data of ascending aortic diameter, Western blotting for Serpine1, and collagen deposition in human TAAs did not pass Shapiro-Wilk or Brown-Forsythe tests, Log transformation was applied for these data. After confirming homogeneous variances and normality, two-group or multi-group comparisons for means were performed by two-sided Student’s t-test or two-way analysis of variance (ANOVA) with Holm-Sidak multiple comparison test, respectively. The log-rank test was used to compare the incidence of ascending aortic rupture between groups. Other causes of death, such as abdominal aortic rupture, were treated as censored events. P<0.05 was considered statistically significant. Statistical analyses mentioned above were performed using SigmaPlot version 14.0 (SYSTAT Software Inc.).

Statistical analyses for proteomic data were performed with R (version 4.1.0).^17^ Proteins in the principal component analysis (PCA) plot were clustered by k-means algorithm (k=2). Differentiated proteins, transformed by “log1p” function in R, between saline and AngII were identified with a Student’s t-test based on the false discovery rate (FDR) adjusted P-value (Q-value) with less than 0.05 as a threshold. Enrichment analyses for gene ontology (biological process) and Kyoto Encyclopedia of Genes and Genomes (KEGG) were performed using “clusterProfiler (4.0.2)” R package.^18^ Heatmaps for comparisons in the standardized protein abundances (i.e. Z scores) between saline vs AngII were generated for differentiated proteins related to SMC contraction, matrix metalloproteinase (MMP), TGFβ signaling, extracellular matrix (ECM)-related molecules. Protein-protein interaction networks were examined using the STRING database (version 11.0) in Cytoscape (version 3.8.0).^19^ Interactions were acquired using the following thresholds: confidence interaction scores≥0.4, active interaction sources=text mining, experiments, databases, co-expression, neighborhood, gene fusion, co-occurrence.

scRNAseq data analyses were also performed with R (version 4.1.0).^17^ Seurat package was used to identify cell types.^20^ Two Seurat objects for each of the mapped unique molecular identifier (UMI) count datasets from mGFP positive cells (Ctrl-SHF and AngII-SHF) were built using the “CreateSeuratObject” function with the following criteria: ≥3 cells and ≥200 detected genes. Cells expressing less than 200 or more than 5,000 genes were filtered out for exclusion of non-cell or cell aggregates, respectively. Cells with more than 10% mitochondrial genes were also excluded. Each of the UMI counts was then normalized as the following: counts for each cell were divided by total counts, multiplied by 10,000, and transformed to natural-log. “FindIntegrationAnchors” and “IntegrateData” functions were used to remove batch effects and integrate the two normalized UMI count datasets. Uniform manifold approximation and projection (UMAP) dimensionality reduction to 20 dimensions for the first 30 principal components (PCs) with 0.1 or 0.5 resolution was applied to identify cell clusters using the normalized- and scaled-UMI count data. “FindAllMarkers” and “FindConcervedMarkers” functions were used to identify conserved marker genes to determine cell types of each of the clusters. The difference of cellular proportions between infusions were examined by Chi-square test. Using “zinbwave (1.14.1)” and “edgeR (3.34.0)” R packages,^21, 22^ differentially expressed genes (DEGs) were identified with a zero-inflated negative binomial regression model based on Q-value<0.05 as a threshold. Gene ontology enrichment analyses in scRNAseq was performed using “clusterProfiler (4.0.2)” in genes with Q-value<0.05 and |Log_2_fold change|>0.5. Monocle package was used for trajectory pseudotime analysis.^23^

Cell-cell interaction was evaluated using a list containing 2,009 ligand-receptor pairs, developed by Skelly et al.^24^ The presence of ligand-receptor pairs within and between each cell type was defined if more than 20% of cells in that cell type had non-zero read counts for the gene encoding ligand or receptor. Then, two cell types were linked and network plots were generated using “igraph” package on R.^25^

For human scRNAseq analyses, read count data were downloaded from Gene Expression Omnibus (GSE155468).^26^ TAA samples were harvested from the ascending region of patients who underwent a surgery for aortic replacement (n=8, mean age=56-78 years, 4 males, 4 females, mean aortic diameters= 4.9-5.8 cm) and control samples were obtained from recipients of heart transplants or lung donors (n=3, mean age=61-63 years, 1 male, 2 females). mRNA abundance was evaluated in SMC and FB clusters. DEG analyses in SMCs and FBs were performed on R using the Hurdle model adjusted for age implemented in the MAST R package.^27^

## Results

### Medial gradients of aortic pathologies in patients with TAAs and AngII-infused mice

We first performed histological analyses to determine the pathological patterns in human sporadic TAA tissues. Immunostaining for α-smooth muscle actin and Movat’s pentachrome staining were performed to detect SMCs and ECM, respectively. Immunostaining for α-smooth muscle actin revealed that the outer media had significantly lower positive areas for α-smooth muscle actin than the inner media in human TAA tissues (**Figure 1A**). Movat’s pentachrome staining identified higher collagen deposition in the outer media (**Figure 1B**).

**Figure 1.**
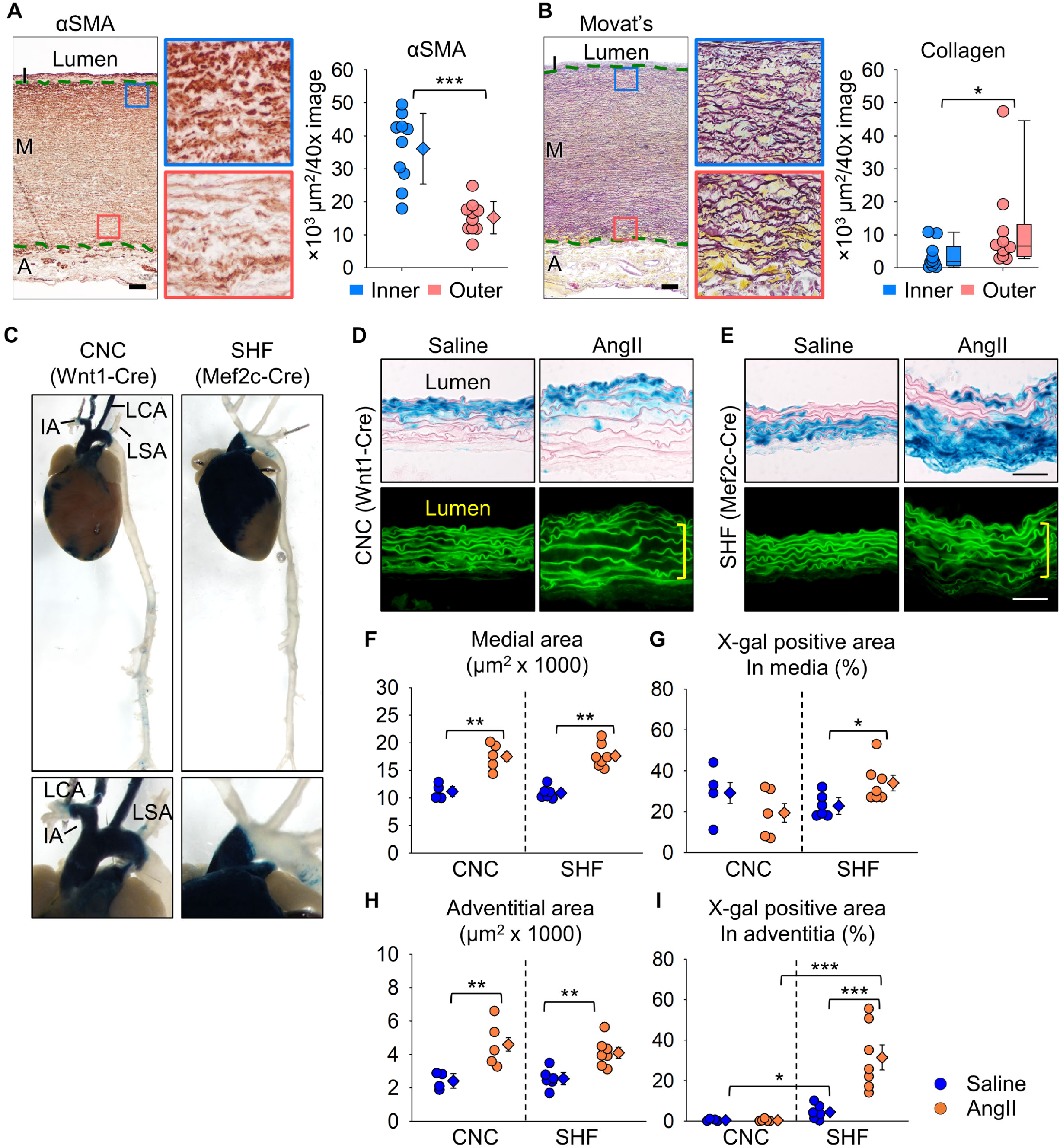
Transmural gradients of aortic pathologies in patients with sporadic TAAs and AngII-infused mice. Representative images of **(A)** immunostaining for α-smooth muscle actin (red) and **(B)** Movat’s pentachrome staining (elastin: dark purple, collagen: yellow). High magnification images are captured from the inner (blue box) and outer (red box) media (n=10). I indicates intima; M, media; A, adventitia. Green dotted lines indicate internal and external elastic layers. αSMA positive and collagen deposited areas were quantified in inner and outer media. Log transformation was applied for the data of collagen deposition. **(C)** Representative images of X-gal stained (blue) aortas from Wnt1- and Mef2c-*Cre* ROSA26R*^LacZ^* mice. IA indicates innominate artery; LCA, left common carotid artery; LSA, left subclavian artery. High magnification images of the proximal thoracic aortas are captured. Representative X-gal staining and FITC images of aortas from either saline- or AngII-infused **(D)** Wnt1- and **(E)** Mef2c-Cre ROSA26R*^LacZ^* mice (n=5 to 7 per group). Yellow square brackets indicate medial thickening. Scale bar, 50 µm. Dot plots for **(F)** medial and **(H)** adventitial areas and X-gal positive area in the **(G)** media and **(I)** adventitia. Diamonds and error bars indicate the mean and SEM, respectively. *P<0.05, **P<0.01, ***P<0.001 by Student’s t-test or two-way ANOVA followed by Holm-Sidak test.

A cell lineage study was subsequently performed to investigate the interaction of SMC origins to medial pathologies. CNC- and SHF-derived cells were tracked using Wnt1- or Mef2c-*Cre* ROSA26R*^LacZ^* mice infused with AngII, respectively. Consistent with our previous study,^13^ the ascending aorta was composed of both CNC- and SHF-derived SMCs (**Figure 1C**). SHF-derived cells were restricted to the ascending aorta, while CNC-derived cells extended into the aortic arch. Although not altering distribution of cells of SMC origins (**Figure 1D, E**), AngII increased medial area (**Figure 1F**). Of interest, X-gal positive areas were increased significantly in medial SHF-, but not CNC-, derived cells (**Figure 1G**). Because selected adventitial FBs are derived from the SHF,^13^ we also evaluated the effects of AngII infusion on cells in the adventitia. Similar to the media, the adventitial area was also increased by AngII infusion (**Figure 1H**) and increased X-gal positive areas were observed in SHF-derived, but not CNC-derived, cells (**Figure 1I**). Thus, TAAs have a transmural gradient of aortic pathologies in humans and mice. The linear tracing data support the notion that SHF-derived cells are susceptible to AngII-induced outer medial and adventitial thickenings.

### AngII altered aortic protein profiles related to extracellular matrix organization

To uncover the molecular mechanisms of AngII-induced ascending aortic pathologies, we performed mass spectrometry-assisted proteomics of the ascending aorta of AngII-infused mice. Ascending aortic tissues were harvested at day 3 of either saline or AngII infusion. Aortic samples showing intramural hemorrhage or dissection were excluded. Proteomic analysis identified 2,644 proteins among all aortic samples. In PCA using the unfiltered proteome, AngII infusion altered protein profiles significantly (45% variation along PC1, **Figure 2A**). Compared to saline-infused mice, AngII infusion changed 596 protein abundances (**Figure 2B**, up- and downregulated: 384 and 212 proteins, respectively). Gene ontology and KEGG pathway enrichment analyses were also performed using these differentially expressed proteins. In addition to terms related to translation and ribosome, ECM related terms, such as “Extracellular matrix organization” and “Proteoglycan in cancer”, were identified in both gene ontology and KEGG analyses (**Figure 2C, Supplemental Figure IA**). A protein-protein interaction analysis also demonstrated the presence of ECM-related molecules in the differentially expressed proteins (**Supplemental Figure IB**). Therefore, we assessed ECM-related molecules comprehensively (**Figure 2D**). Several contractile proteins, including Myh11, were downregulated by AngII, while Mmp2 (matrix metalloproteinase 2) was upregulated in AngII-infused mouse aortas. In TGFβ signaling molecules, AngII upregulated Ltbp2 (latent TGFβ binding protein 2) and Tgfb2 (TGFβ-2). AngII also increased several ECM components, Col3a1 (collagen type III alpha 1 chain), Col5a2 (collagen type V alpha 2 chain), and Eln (elastin). Other ECM-related molecules, Thbs1 (thrombospondin 1), Fn1 (fibronectin 1) and Tnc (tenascin C), were also upregulated by AngII infusion. Thus, during the short interval of 3 days of infusion, AngII changed ECM-related proteome significantly in the ascending aorta.

**Figure 2.**
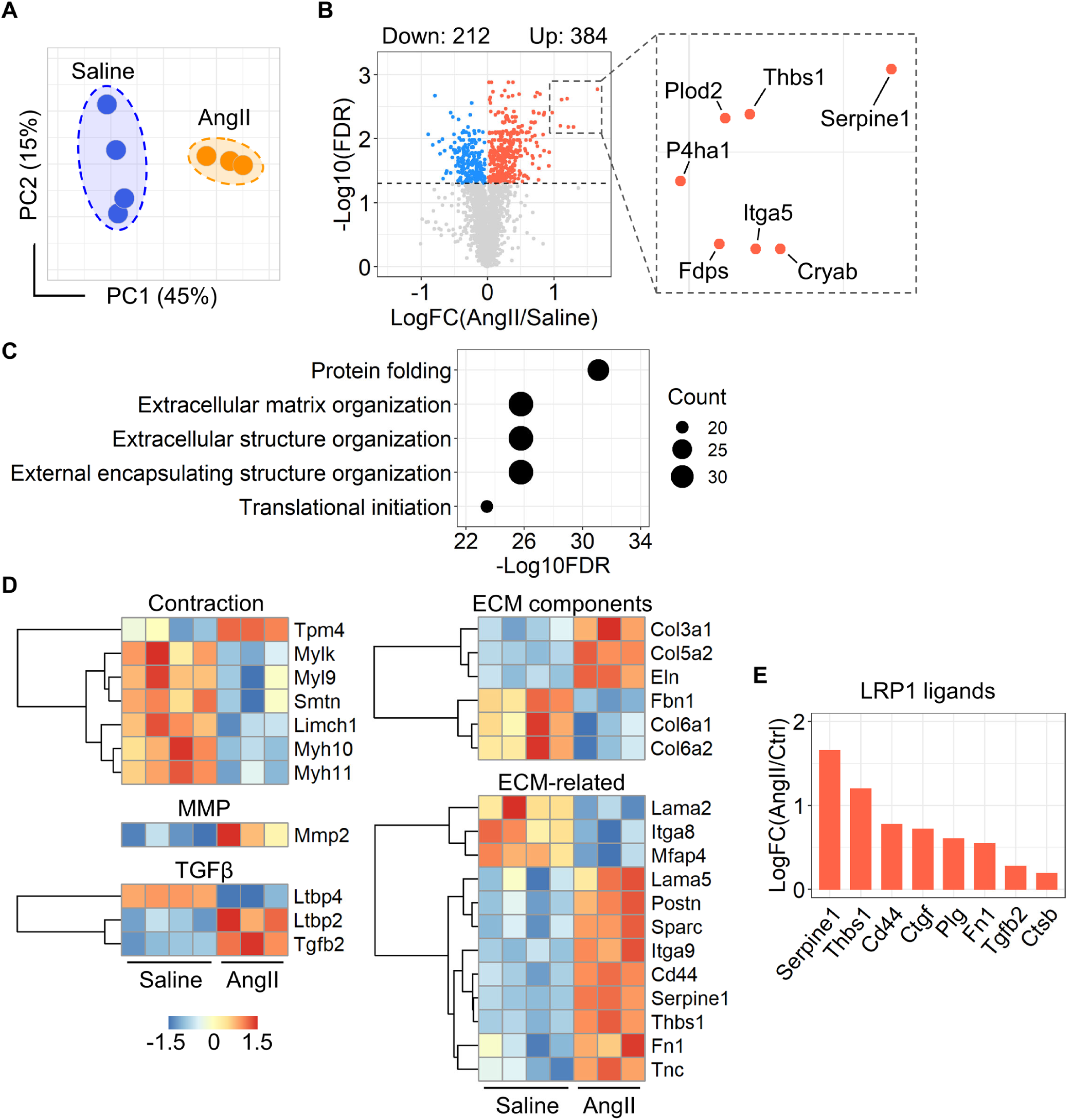
AngII altered proteins related to ECM in the ascending aorta of mice. **(A)** Principal component analysis of the unfiltered proteome. PC1/PC2 indicate first/second principal components (n=3 or 4 per group). **(B)** Volcano plots for differentiated proteins between in ascending aortas of saline and AngII-infused mice. **(C)** Top 5 annotations of enrichment analyses in gene ontology - biological process. **(D)** Heatmap with Z score coloring for proteins related to SMC contraction, MMP, TGFβ signaling, ECM-related molecules compared between saline and AngII infusion. **(E)** Log fold change of LRP1 ligands.

Low-density lipoprotein receptor-related protein 1 (LRP1) is a multifunctional protein that plays a role in ECM maturation.^28^ LRP1 deletion in pan SMCs causes aortic pathologies including aneurysm and tortuosity with ECM degradation.^29–32^ Thus, we examined protein abundances of LRP1 and its ligands. Aortic LRP1 protein abundance was not changed by AngII infusion (data not shown, P=0.89); however, multiple LRP1 ligands, including Serpine1 (plasminogen activator inhibitor 1, PAI1), were increased by AngII (**Figure 2E**). Of note, a volcano plot demonstrates that Serpine1 was the most significantly increased molecule by AngII infusion (**Figure 2B**). Since these ligands are internalized into the cytosol through LRP1 for their degradation,^28^ these data suggest the impairment of aortic LRP1 function. Taken together, the aortic proteome related to ECM was altered as early as day 3 by AngII infusion with the increase of multiple LRP1 ligands.

### LRP1 deletion in SHF-derived cells augmented AngII-induced thoracic aortopathies

As described above, our proteomic analysis, in combination with our previous findings that deletion of LRP1 in pan SMCs augmented TAA formation with ECM degradation in AngII-infused mice,^30^ prompted us to determine whether LRP1 deletion in SHF lineage affects AngII-induced TAA formation. LRP1 was deleted in SHF-derived cells using a Mef2c promoter driven *Cre*. In Mef2c-*Cre*-expressing mice, *Cre* activity was observed mainly in the outer media (**Supplemental Figure IIA**), and the *Lrp1 delta flox* PCR fragment was detected in addition to a band of native *Lrp1* (**Supplemental Figure IIB-D**), confirming DNA recombination of LRP1 in SHF-derived cells. LRP1 protein abundance was reduced by 37% in the ascending aortic media of SHF-specific LRP1 deleted mice compared to *Cre* negative littermates with LRP1 deletion restricted to the outer media of the ascending aorta (**Supplemental Figure IIE-F**). Mass spectrometry-assisted proteomics also verified reduction of LRP1 peptide intensity in the ascending aorta of Mef2c-*Cre* expressing mice (**Supplemental Figure IIG**).

SHF-specific LRP1 deletion did not affect growth or development of mice. To promote TAAs, AngII was infused subcutaneously into mice for 4 weeks. LRP1 deletion in SHF-derived cells augmented AngII-induced aortic dilation in the ascending aorta as demonstrated by both ultrasound and in situ imaging (**Figure 3A, B**). SHF-specific LRP1 deletion also exacerbated AngII-induced ascending aortic rupture and elastin fragmentation (**Figure 3C-E**). Systolic blood pressure was not different between genotypes in response to AngII infusion (**Figure 3F**), demonstrating that augmented aortopathy was independent of systolic blood pressure. These data support that SHF-derived cells exert a critical role in the structural integrity of the ascending aorta.

**Figure 3.**
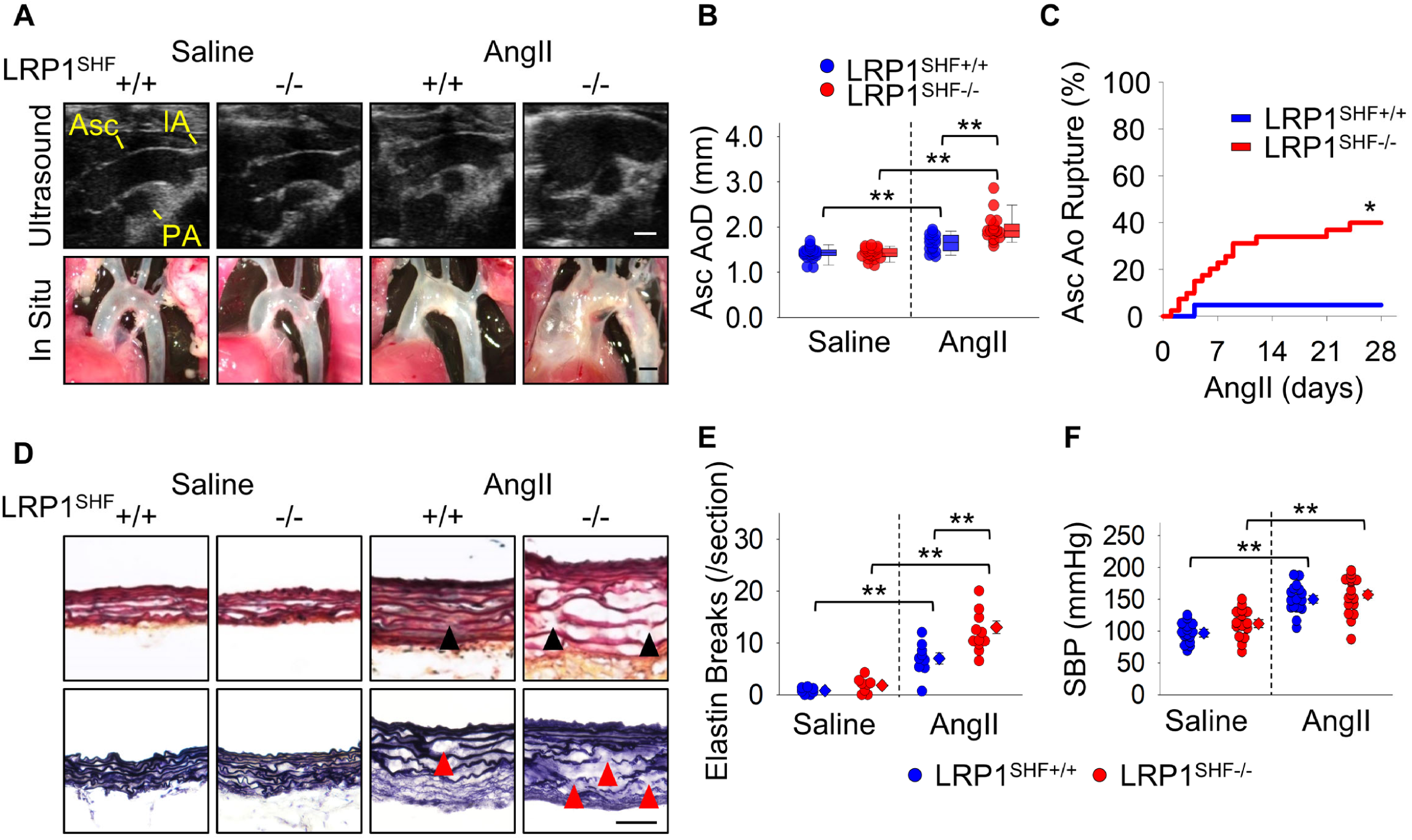
LRP1 deletion in SHF-derived cells augmented AngII-induced TAA formation and aortic rupture in mice. **(A)** Representative ultrasound images and in situ gross appearances of proximal thoracic aortas (n=21 to 40 per group). Asc indicates ascending aorta; IA, innominate artery; PA, pulmonary artery. Scale bar, 1 mm. **(B)** Ascending aortic diameter (Asc AoD) measured by ultrasonography. **(C)** Cumulative incidence of ascending aortic (Asc Ao) rupture. **(D)** Representative Movat’s and Verhoeff iron hematoxylin staining. Scale bar, 50 µm. Arrow heads indicate elastin fragmentation. **(E)** Elastin break counts in Movat’s stained sections. **(F)** Systolic blood pressure. Diamonds and error bars indicate the mean and SEM, respectively. *P<0.01 by log-rank test, **P<0.001 by two-way ANOVA followed by Holm-Sidak test. Log transformation was applied for the data of ascending aortic diameters.

### AngII infusion compromised TGFβ signaling in SHF-derived SMCs and FBs

Since using tissues of fate mapping mice enables unbiased transcriptomics profiling in a lineage-specific manner, we investigated molecular roles of SHF-derived cells in AngII-induced TAA formation by scRNAseq in aortas from Mef2c-*Cre* ROSA26R*^mT/mG^* mice. Aortic tissues were harvested from Mef2c-*Cre* ROSA26R*^mT/mG^* male mice at baseline and after 3 days of AngII infusion. Four to five aortic tissues were pooled to obtain sufficient number of aortic cells (**Figure 4A**). Since mGFP protein was present on Mef2c-*Cre* expressing cells of this mouse strain, mGFP positive cells were derived from the SHF (**Figure 4B**). After sorting cells based on mGFP signal, scRNAseq was performed using mGFP positive cells.

**Figure 4.**
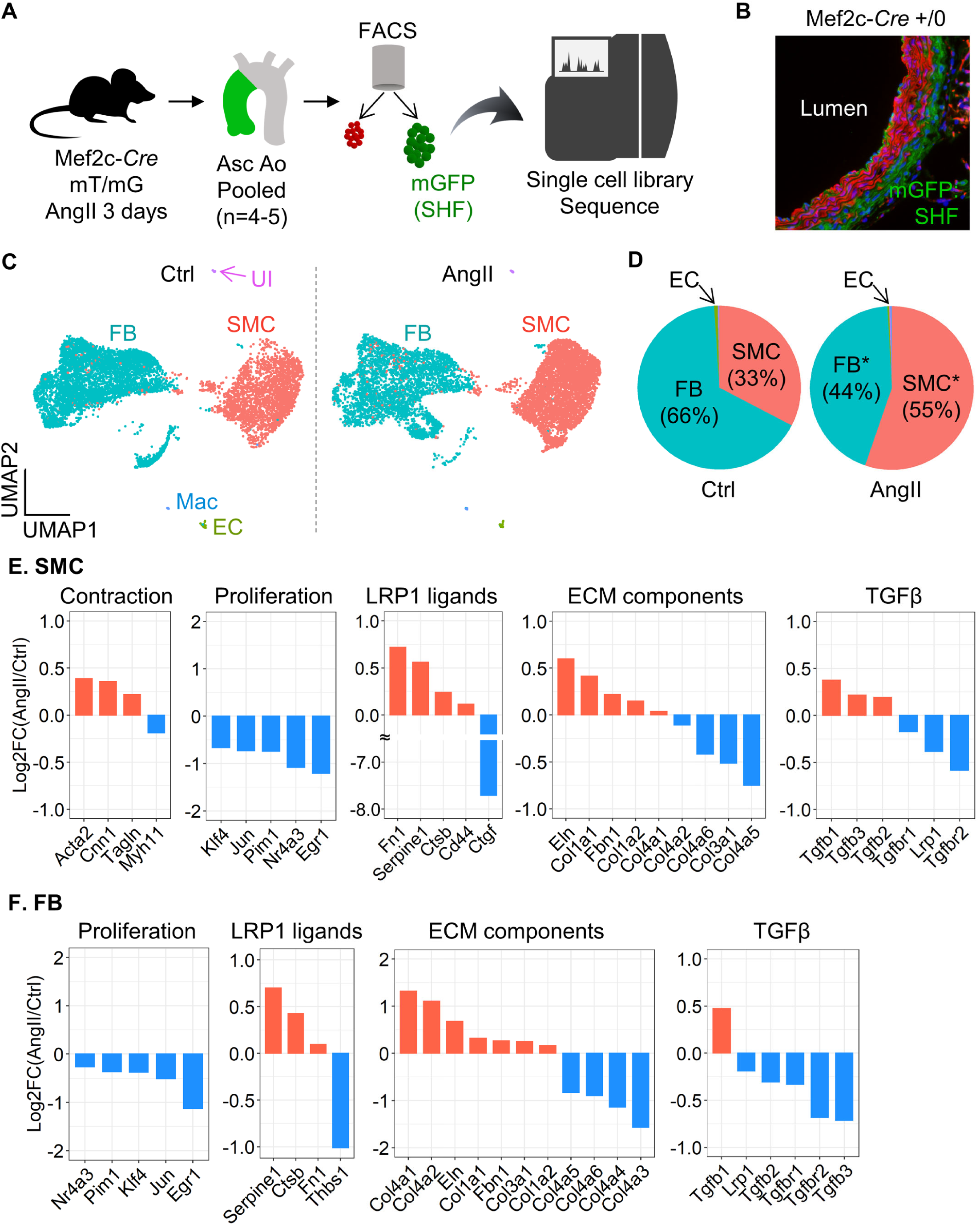
Significant reduction of *Tgfbr2* mRNA in SHF-derived SMCs and FBs of AngII-infused mice. **(A)** Experimental design of scRNAseq using Mef2c-*Cre* +/0 ROSA26R*^mT/mG^* mice. **(B)** Representative fluorescent image of the ascending aorta from Mef2c-*Cre* +/0 ROSA26R*^mT/mG^* mice. **(C)** UMAP plots of aortic cells at baseline (Ctrl) and after 3 days of AngII infusion. SMC indicates smooth muscle cells; FB, fibroblasts; EC, endothelial cells; Mac, macrophages; UI, unidentified cells. **(D)** Pie charts for the composition of each cell type. *P<0.0001 by Chi-square test. Log_2_ fold change of mRNA abundance related to SMC contraction, cell proliferation, LRP1 ligands, ECM components, and TGFβ ligands and receptors between Ctrl vs AngII in **(E)** SMCs and **(F)** FB clusters.

The two normalized datasets were properly integrated (**Figure 4C**). scRNAseq analysis revealed multiple cell types, including SMC, FB, and endothelial cells, in the mGFP cell population (**Figure 4C, D, Supplemental Figure IIIA, B**). At baseline, SHF-derived SMCs were composed of 33% SHF-derived cells and it was increased to 55% by AngII infusion (**Figure 4D**). Meanwhile, FB population was decreased from 66% to 44%. Since SMCs and FBs were comprised the majority, DEG analysis was performed in these clusters. AngII altered 3,848 (up: 1,862; down: 1,986) and 3,398 (up: 1,931; down: 1,467) genes in SMCs and FBs, respectively (**Supplemental Figure IVA, VA**). Although the top 10 up- and downregulated genes were variable between SMCs and FBs, except for *Ptrf* (**Supplemental Figure IVB, VB**), enrichment analysis identified “extracellular matrix organization” as a common term in SMC and FB clusters (**Supplemental Figure IVC, VC**). Therefore, similar to the proteomic analyses, we next examined ECM-related molecules in the scRNAseq data (**Figure 4E, F**). In SMCs, *Acta2*, *Cnn1*, and *Tagln*, contractile genes, were increased after 3 days of AngII infusion. Of note, proliferation genes, such as *Klf4* and *Egr1*, were downregulated in both SMCs and FBs by AngII infusion. As LRP1 ligands, *Serpine1, Ctsb* (cathepsin B), and *Fn1* (fibronectin 1) were increased by AngII in SHF-derived SMCs and FBs. Multiple ECM components, *Eln*, *Col1a1*, *Col1a2*, and *Col4a1*, were increased in both SMC and FB clusters in response to AngII infusion. However, TGFβ ligands demonstrated variable responses to AngII between cell types. AngII increased *Tgfb1-3* in SMCs, but *Tgfb2-3* were decreased in FBs. It is noteworthy that TGFβ receptors, including *Lrp1* and *Tgfbr2*, were significantly downregulated by AngII in both SMCs and FBs.

mRNA abundance of TGFβ ligands and receptors were also examined in human scRNAseq data (GSE155468).^26^ SMCs and FBs were extracted and mRNA abundance of *TGFB1-3*, *TGFBR1-2,* and *LRP1* were compared between control and TAA samples. In SMCs, TGFβ ligands and receptors, including *LRP1*, were decreased modestly in TAAs compared to control aortas (**Supplemental Figure VIA**). In contrast, TGFβ ligands and receptors were increased in FBs of TAAs, except for *TGFB1* (**Supplemental Figure VIB**).

### A unique sub-cluster of SHF-derived fibroblasts in ascending aortas of AngII-infused mice

Since the aortic wall comprises mostly SMCs and FBs (**Figure 4C, D**), we performed further sub-clustering analyses in SHF-derived SMCs and FBs. SHF-derived SMCs had 3 distinct clusters, but there was no remarkable alteration during AngII infusion in transcriptomic distributions of these cells (**Supplemental Figure VII**). Meanwhile, a unique sub-cluster was observed in SHF-derived FBs of AngII-infused mice (FB4, **Figure 5A**). The FB4 sub-cluster was composed of 10 and 544 cells at baseline and after 3 days of AngII infusion, respectively (**Supplemental Figure VIIIA**). AngII infusion altered 586 genes in the FB4 sub-cluster and these genes were significantly associated with cytokinesis, as identified by gene ontology analysis (**Supplemental Figure VIIIB, C**). To further characterize the FB4 sub-cluster, featured genes were determined by comparing the transcriptomes among all FB sub-clusters. Genes related to cell division, such as *H2afz*, *Cks2*, *Cenpa*, *Cdk1*, and *Mki67*, were identified uniquely in the FB4 sub-cluster (**Figure 5B, Supplemental Figure IX**). These results indicate that FB4 cells had enhanced proliferative features. However, FB4 cells were not highly abundant for *Ly6a*, as a marker for Sca1+ FBs (**Figure 5C**). While the transcriptome of FB4 differed from other FB sub-clusters (**Figure 5D**), the ligand-receptor interaction analyses in SMC and FB clusters demonstrated accelerated cell-cell interactions among cell types by AngII infusion (**Figure 5E**). In particular, FB4 sub-cluster had more apparent interactions with SMC and FB1 clusters under AngII infusion. Trajectory analysis demonstrated that FB4 cells were derived from FB1 cells in a pseudotime (**Figure 5F**). Because of these interactions, we subsequently compared DEGs related to ECM maturation between FB1 and FB4 sub-clusters (**Figure 5G**). LRP1 ligands, including *Serpine1*, were all downregulated in FB4. ECM component genes, such as *Eln* and *Col1a1*, were also lower in FB4. TGFβ receptors were further downregulated in FB4 by AngII infusion. Alongside the scRNAseq analyses in SMCs and FBs, these data suggest that AngII compromises the TGFβ signaling pathway in SHF-derived cells, which may contribute to the susceptibility of SHF-derived cells to vascular pathologies.

**Figure 5.**
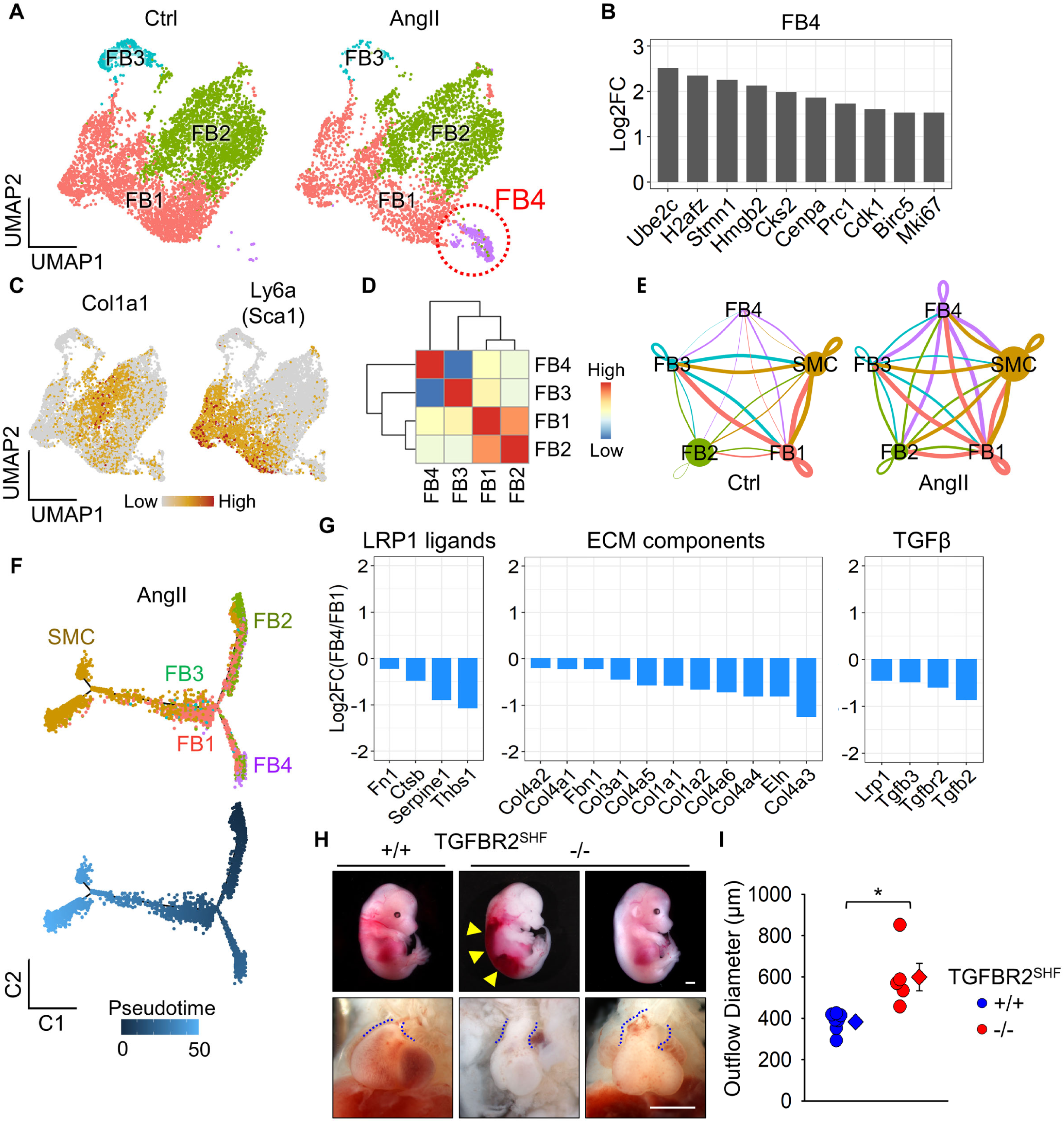
A distinct sub-cluster with decreased *Tgfbr2* mRNA in SHF-derived fibroblasts of AngII-infused mice. **(A)** UMAP plots at baseline (Ctrl) and after 3 days of AngII infusion in SHF-derived FBs. **(B)** Log_2_ fold change of mRNA abundance for top 10 unique genes in FB4 sub-cluster. **(C)** Featured plots for *Col1a1* and *Ly6a* in FB sub-clusters. **(D)** Heatmap for cluster similarity among FB sub-clusters. **(E)** Ligand-receptor interactions among aortic cells. Lines connect to cell populations that express cognate ligands or receptors. The line thickness and node size correspond to the number of cognate ligand-receptor expression and cell number, respectively. **(F)** Trajectory analysis in SMC and FB sub-clusters from AngII-infused mice. **(G)** Log_2_ fold change of mRNA abundance related to LRP1 ligands, ECM components, and TGFβ molecules between FB1 and FB4 sub-clusters after AngII infusion. **(H)** Representative images of gross appearance and the outflow tract of wild type littermates (left), dead (middle), and survived (right) fetuses with SHF-specific TGFBR2 deletion at E12.5. Blue dotted lines indicate the edge of the outflow tract. Yellow triangles indicate retroperitoneal hemorrhage. **(I)** Outflow diameter was measured at 300 to 400 µm distal to the root of viable fetuses at termination (n=5 to 7 per group). Scale bar, 1 mm. Diamonds and error bars indicate the mean and SEM, respectively. *P<0.001 by two-tailed Student’s t-test.

To investigate the impact of TGFβ signaling pathway in SHF-derived cells on vascular integrity, we deleted TGFBR2 in SHF-derived cells in mice. Of note, SHF-specific TGFBR2 deletion led to embryonic lethality with retroperitoneal hemorrhage beginning at E12.5 (**Figure 5H**). In addition, the outflow tract was dilated significantly in fetuses with SHF-specific TGFBR2 deletion compared to wild type littermates (**Figure 5H, I**). Thus, TGFBR2 deficiency in SHF-derived cells led to prenatal vasculopathies, suggesting the importance of SHF-derived cells through TGFBR2 for vascular development and maintaining of its integrity.

### Transmedial gradient of PAI1 in ascending aortas of AngII-infused mice and sporadic ascending TAAs in humans

We integrated DEG results of proteomics and scRNAseq data from mice. There were 412 overlapped genes between proteomics and scRNAseq using the SHF-SMC cluster, and 198 molecules were upregulated in both datasets (highlighted in red, **Figure 6A**). In SHF-FBs, 375 genes were overlapped, and 205 genes were increased (**Figure 6B**). As LRP1 ligands, Serpine1, Fn1, and Ctsb were consistently greater abundance in AngII-infused mice, as shown in proteomics and scRNAseq analyses of SMCs and FBs (**Figure 6A, B**). In protein analyses, PAI1 (*Serpine1*) was the most significantly increased molecule by AngII (**Figure 6A, B**). Western blot analysis showed a striking increase (∼26 fold) of PAI1 in ascending aortas from mice infused with AngII (**Figure 6C**). Immunostaining of PAI1 was hardly detected in aortas from control mice, whereas it was intense in ascending aortas of AngII-infused mice (**Figure 6D**). In ascending aortas of AngII infused mice, PAI1 was present through the aortic wall; aortic PAI1 was dominant in the outer media and the adventitia, coincident with the gradient of medial pathologies (**Figure 6D**).

**Figure 6.**
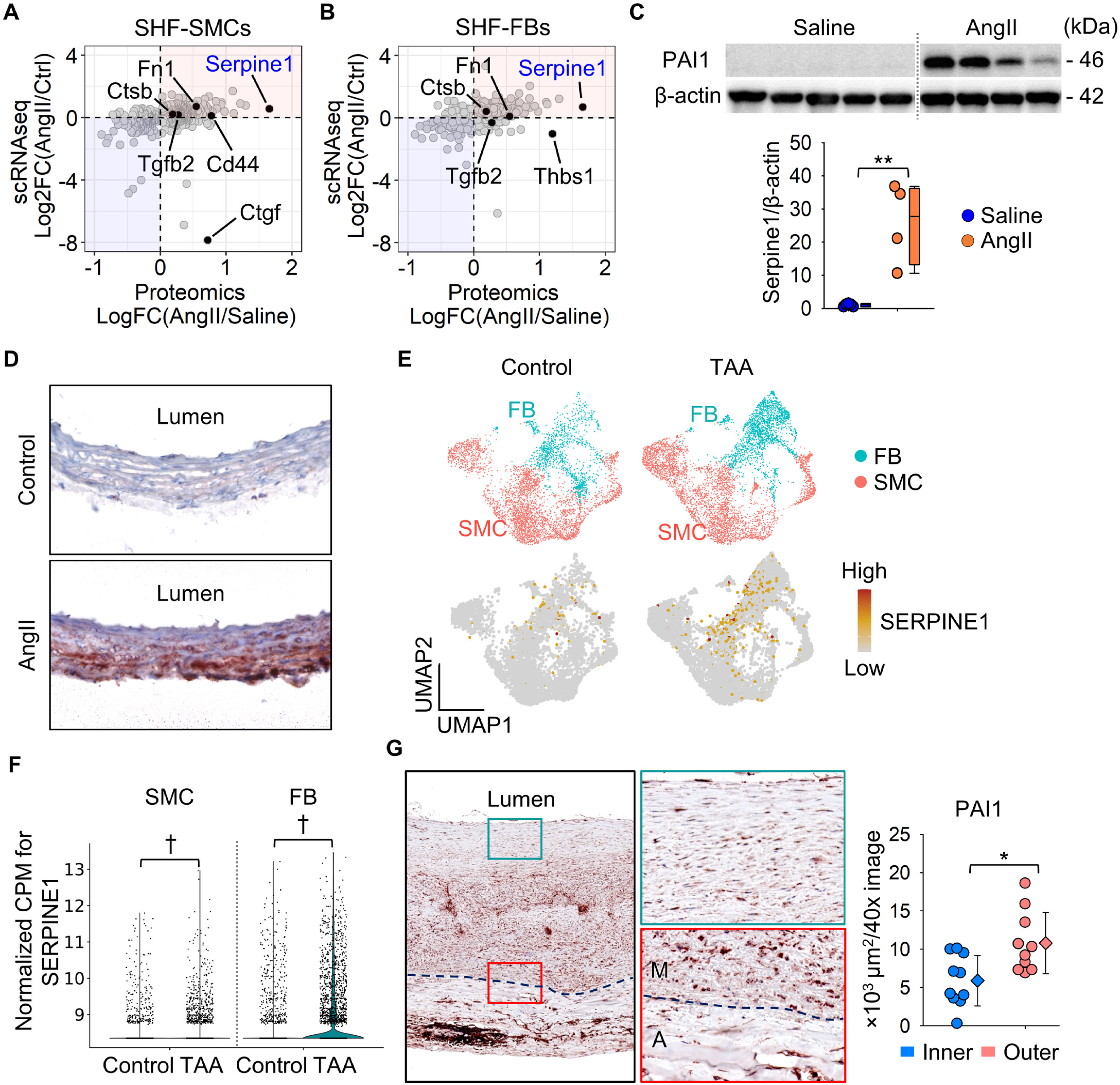
Integration of proteomic and scRNAseq results identified PAI1 (*Serpine1*) as a key contributor to the medial gradient of aortic pathologies. Scatter plots for Log or Log_2_ fold change of protein and mRNA abundances. Data were compared between proteomics and scRNAseq in SHF-derived (A) SMC and (B) FB clusters. Selected LRP1 ligands were highlighted. (C) Western blot analysis for PAI1 (Serpine1) and β-actin in the whole ascending aorta from saline or AngII-infused mice (n=4-5 per group). **P<0.001 by Student’s t-test after Log transformation. (D) Representative images of immunostaining for PAI1 (red) in ascending aortas from control or AngII-infused mice (n=7 per group). (E) UMAP and (F) violin plots for *SERPINE1* in SMC and FB clusters of control subjects and patients with ascending TAAs (n=3 and 8 per group, scRNAseq data were downloaded from GSE155468). †P<0.001 analyzed using the Hurdle model adjusted for age implemented. (G) Representative images of immunostaining for PAI1 in patients with sporadic ascending TAA (n=10) and its quantification. Dotted lines indicate external elastic laminae; M, media; A, adventitia. Diamonds and error bars indicate the mean and SEM, respectively. *P<0.01 by Student’s t-test.

For clinical relevance, we evaluated *SERPINE1* mRNA abundance and protein distribution in human aortic samples. In the human scRNAseq data (GSE155468),^26^ *SERPINE1* abundance was assessed in SMC and FB clusters. UMAP plots showed the presence of *SERPINE1* in both SMC and FB clusters of control and TAA samples (**Figure 6E**); *SERPINE1* was significantly higher in SMCs and FBs of human TAA samples compared to controls (**Figure 6F**). Immunostaining demonstrated that PAI1 was expressed in the aortic media and adventitia of human TAA samples (**Figure 6G**). Consistent with AngII-infused mouse aortas (**Figure 6D**), highly abundant areas of PAI1 were observed in the outer media adjacent to the external elastic lamina in most TAA tissues (**Figure 6G**).

## Discussion

Our study demonstrated that (1) aortic pathologies were mainly observed in the outer media of the ascending aorta in patients with TAAs and AngII-infused mice, (2) SHF-derived cells were responsible for AngII-induced medial and adventitial thickening in the ascending aorta, (3) AngII altered protein profiles related to ECM and LRP1 ligands, (4) AngII decreased *Tgfbr2* mRNA in SHF-derived SMCs and FBs, (5) AngII led to a distinct sub-cluster of SHF-derived FBs that had lower *Tgfbr2* mRNA abundance, (6) Either LRP1 or TGFBR2 deletion in SHF-derived cells led to vasculopathies in mice, and (7) proteomic and scRNAseq analyses identified PAI1 as a key molecule that showed a transmedial gradient in the ascending aorta of AngII-infused mice or human TAA tissues. Collectively, this study provides multiple evidence to support the importance of SHF-derived cells in maintaining the integrity of aortic wall.

A number of studies have shown an important role for TGFβ signaling in maintaining aortic wall integrity.^9, 33–35^ Both LRP1 and TGFBR2 are related to the TGFβ signaling pathways,^28, 33^ and genetic deletions of these molecules in pan-SMCs promotes TAA formation in mice.^8, 9, 29–31, 36^ Therefore, in the present study, the biological effects of SHF-derived cells in aortic integrity was investigated by SHF-specific deletion of either LRP1 or TGFBR2. Of note, SHF-specific TGFBR2 deletion led to vascular malformation and SHF-specific LRP1 deletion recapitulated aortic phenotypes in mice with its deletion in pan-SMCs. In the ascending aorta, SHF-derived cells comprise approximately half of SMCs and selected FBs.^13^ Thus, we concluded that SHF-derived cells were functionally important to maintain the structural integrity of vascular wall.

The present study revealed that deletion of either TGFBR2 or LRP1 in SHF-derived cells promotes vasculopathies. These data are consistent with the outer layers of SMCs derived cells being crucial for development of vascular pathologies related to TGFβ signaling. A previous study using mice expressing a fibrillin1 mutation (*Fbn1*^C1041G/+^) reported that heterozygous deletion of *Notch1* in SHF-derived cells had a trend for augmenting luminal dilatation of the aortic sinus.^37^ In contrast, another study demonstrated that Smad2 deletion in CNC-derived cells reduced dilation of the aortic sinus in a mouse model of Loeys-Dietz syndrome.^38^ Since there are aortic region-specific responses to TGFβ,^39^ these two developmental origins may have divergent pathophysiological functions in different aortic regions.

Elastin fragmentation is a pathological feature of TAAs and is associated with decrease of aortic resilience.^40, 41^ Elastic fibers are one of the major matrix components and vital for maintaining aortic structure.^42^ Mutations of the elastin gene cause aortic stenosis due to overgrowth of SMCs.^40, 41^ Also, SMC-specific elastin deletion induces aortic extension and coarctation in mice.^43^ Our proteomic and scRNAseq analyses demonstrated increased aortic elastin in response to AngII infusion. These data suggest that de novo elastin synthesis is a compensatory reaction for maintaining aortic integrity during AngII stimulation. It is of note that major TGFβ receptors, *Tgfbr2* and *Lrp1*, were downregulated during AngII infusion for 3 days in SHF-derived SMCs and FBs, although the TGFβ signaling pathway plays an important role in ECM maturation.^44^ Hence, increase of aortic elastin in SHF-derived cells may not be regulated through the TGFβ signaling pathway directly during this interval. In addition, multiple collagen genes showed variable responses to AngII infusion. Therefore, it remains to be clarified which ECM component molecules are regulated during AngII infusion.

Our proteomic and scRNAseq analyses identified PAI1 as a key molecule in AngII-induced TAA formation. PAI1 has been implicated in thoracic aortic diseases by omics approaches in several previous studies.^45, 46^ A microarray of aortas from patients with aortic dissections revealed an increase of PAI1 mRNA abundance in dissected aortas.^45^ scRNAseq using aortas from patients with Marfan syndrome and a mouse model of this disease demonstrated the presence of PAI1 in aortic SMCs.^46^ Consistent with these reports, the present study also identified PAI1 in the ascending aorta shortly after initiation of AngII infusion. Of interest, aortic PAI1 was increased dramatically by AngII. Furthermore, our histological analyses found a transmedial gradient of PAI1 in AngII-infused mice and human TAA samples. The distinct distribution of PAI1 was matched with the medial gradient of aortic pathologies, consistent with PAI1 contributing to the complex mechanism of TAA formation.

Several articles have investigated a role of PAI1 in abdominal aortic aneurysms.^47–49^ Focal overexpression of PAI1 inhibit aneurysm formation in the abdominal aorta of a rat model using xenograft.^47^ Adventitial overexpression of PAI1 also protect abdominal aneurysm formation in AngII-infused mice.^48^ In addition, systemic deletion of PAI1 augmented elastase-induced abdominal aortic aneurysms in mice.^49^ Therefore, PAI1 plays a protective role in development of abdominal aortic aneurysms. However, molecular mechanisms underlying TAAs and abdominal aortic aneurysms are distinct. There is a lack of evidence showing biological function of PAI in thoracic aortopathies. PAI1 is a ligand for LRP1 and involved in the TGFβ signaling pathway.^50–52^ In subsequent studies, we will elucidate the role of PAI1 in SHF-derived cells on TAA formation as well as TGFβ signaling pathway through LRP1.

SHF-specific TGFBR2 deletion led to dilatation of the outflow tract with post peritoneal hemorrhage. Thus, TGFBR2 in SHF-derived cells is essential for vascular development and integrity. However, due to the embryonic lethality by constitutive deletion of TGFBR2 in SMCs, the role of TGFBR2 in SHF-derived cells on thoracic aortopathies remains unclear. The Mef2c gene is not sufficiently expressed in the cardiovascular system at the postnatal phase, especially in normal conditions during the adult phase.^53^ Therefore, it is not feasible to activate Mef2c-*Cre* using the ER^T2^ system in adult mice. Development of research methods are needed to overcome the barrier to postnatal gene manipulation in the SHF origin.

Previous studies have reported that prenatal deletion of TGFBR2 in CNC-derived cells results in cleft palate and a congenital heart defect, persistent truncus arteriosus.^54, 55^ Therefore, there is compelling evidence that CNC-derived cells exert a pivotal role in the integrity of the cardiovascular system.^54, 55^ However, the present study focused on the role of SHF-derived cells in thoracic aortopathies, since scRNAseq data revealed the important role of these cells on the regulation of elastin and TGFβ signaling genes. Furthermore, our mouse studies demonstrated vascular phenotypes by gene manipulation in SHF-derived cells. These results support the notion that SHF-derived cells are crucial in maintaining aortic wall integrity. In fact, a recent study reported that a gene TGFBR1 mutation suppressed muscle contractility in SMCs from the SHF, but not CNC, origin.^56^ However, it is still desirable to directly compare the role of two different SMC origins on formation of thoracic aortopathies.

Our scRNAseq identified a distinct FB sub-cluster, and AngII infusion further decreased *Tgfbr2* mRNA abundance in the distinct FB sub-cluster compared to other FBs. Since TGFBR2 deletion in SHF-derived cell led to vascular malformation, significant reduction of *Tgfbr2* mRNA abundance by AngII in this cluster may exert a deleterious role in the pathophysiology of TAAs. scRNAseq enabled us to evaluate cellular heterogeneity and identify a unique cellular cluster, which is a great advantage of scRNAseq. However, it is still difficult to investigate cluster-specific molecular mechanisms. A technical breakthrough is needed to manipulate specific sub-clusters for understanding its role in the pathophysiology of TAAs.

scRNAseq demonstrated that *Tgfbr2* and *Lrp1* were downregulated by AngII infusion in both SMCs and FBs derived from the SHF of mouse aortas. While in human aortas, *TGFBR2* and *LRP1* were decreased in SMCs of TAAs, but increased in FBs of TAAs. *TGFBR2* and *LRP1* in FBs in human scRNAseq conflict with the mouse scRNAseq results. In the mouse scRNAseq, aortic samples were harvested at day 3 of AngII infusion prior to TAA formation, but human scRNAseq used aortas at an advanced stage of diseases that were acquired during surgical repair. As the role of TGFβ signaling changes depending on disease stage,^57^ this may contribute to the difference of scRNAseq results seen between humans and mice in the present study. In addition, since it was not feasible to discern embryonic origins in the human scRNAseq dataset, the data were analyzed using all SMCs and FBs rather than isolated SHF-derived cells. This may also contribute to the difference seen in these scRNAseq results.

In conclusion, our study highlights the functional importance of SHF-derived cells in maintaining vascular integrity using multiple TAA mouse models. This study provides strong evidence that SHF-derived cells exert a role during development of thoracic aortopathies. Heterogeneity of SHF-derived cells contributes to complex mechanisms of aortopathy formation, which should be taken into consideration when investigating the pathogenesis of thoracic aortopathies.

## Non-standard Abbreviations and Acronyms

AngII: angiotensin II
CNC: cardiac neural crest
ECM: Extracellular matrix
FB: fibroblast
LRP1: low density lipoprotein-related protein 1
PAI1: plasminogen activator inhibitor 1
SHF: second heart field
SMC: smooth muscle cell
TAA: thoracic aortic aneurysm
TGF-β: transforming growth factor beta
Tgfbr2: transforming growth factor beta receptor type 2

## Acknowledgments

Study design: HS, HH, CZ, YL, SM, SAS, JZC, DLR, YHS, MA, MWM, HSL, AD

Implementation of animal experiments: HS, CZ, JJM, DAH, DLR

Histological analyses: HS, MKF, DAH, DLR

Data analyses: HS, YK, DLR, HSL, AD

Proteomics and associated informatics: HS, YK, HH, SM, LHL, SAS

scRNAseq and associated informatics: HS, YK, CZ, YL, YHS

Supervising and data verification: SAS, YHS, SAL, MA, MWM, HSL, AD

Writing the draft manuscript: HS

Editing the manuscript: all authors

## Supplemental Methods

### Mice

Mice were housed in ventilated cages with negative air pressure (Allentown Inc). Mouse housing conditions are described in **Supplemental Table II**. Briefly, Aspen hardwood chips were used as bedding (#7090A, Envigo). Mice were fed a normal laboratory rodent diet (#2918, Envigo) and provided with drinking water from a reverse osmosis system (pH 6.0 - 6.2) ad libitum. Ambient temperature ranged from 68 to 74°F and humidity was 50 to 60%. The room’s light:dark cycle was 14:10 hour. For mice expiring before the study endpoint, necropsy was performed to determine cause of death. Genotypes were confirmed at termination using DNA isolated from tails or brains using Maxwell tail DNA purification kits (#AS1120, Promega) and PCR with primers shown in **Supplemental Table III**.

### Embryonic Study

To investigate aortic malformation during the prenatal phase, fetuses were harvested from pregnant females on either E11.5 or E12.5. The morning after detection of a vaginal plug in mated females was defined as E0.5 of gestation. Gravid female mice were euthanized using a ketamine/xylazine cocktail (90 and 10 mg/kg, respectively), and saline (8 ml) was perfused into the left ventricle. The abdominal cavity of females was opened and fetuses were dissected free. Gross appearance of fetuses was recorded using a dissection microscope with a high-resolution camera (#SMZ800, #DS-Ri1, Nikon) and cranial tissue was retrieved for genotyping. Embryos were immersed in buffered formalin (10% wt/vol). Twenty-four hours later, chest wall, pericardium, and atriums were removed gently, and gross appearance of the outflow tract was imaged using a dissection microscope coupled to a high-resolution camera. Diameters of the outflow tract were measured at 300-400 µm distal to the aortic root in viable fetuses at termination.

### Aortic Tissue Processing for Western Blot and Histological Analyses

Aortic tissue was harvested after either 3 days or 4 weeks of infusion for Western blot and histological analyses. Mice were euthanized using a ketamine/xylazine cocktail. The thoracic cavity was cut open and saline (8 ml) was perfused through the left ventricle. For Western blot analysis, aortic samples were harvested and snap-frozen in liquid nitrogen. For image acquisition, a black thin plastic sheet was placed beneath the heart and ascending aorta in situ. Hearts and thoracic aortas were dissected free, and immersed either in paraformaldehyde (PFA, 4% wt/vol) for gross tissue histology or placed in OCT for sectioning. For frozen sectioning, serial cross-sections (10 μm) were collected starting at aortic valves and ending at the innominate artery.

### Histological Analyses

To detect β-galactosidase activity, whole tissues and fresh frozen sections were fixed with PFA for either 1 hour at 4°C or 10 minutes at room temperature, respectively. PFA-fixed tissues were incubated in buffer containing sodium phosphate (100 mM, pH 7.3), MgCl_2_ (2 mM), sodium deoxycholate (0.01% wt/vol) and NP40 (0.02% wt/vol). X-gal (1 mg/ml, V394A, Promega), potassium ferricyanide (5 mM), and potassium ferrocyanide (5 mM) were added to buffer and samples were incubated overnight at room temperature. Whole tissues were subsequently immersed in formalin (10% wt/vol). Tissue sections on slides were rinsed to remove X-gal, and incubated with eosin (1% wt/vol) for 2 minutes, and coverslipped using glycerol gelatin (GG1, MilliporeSigma). Movat’s pentachrome and Verhoeff’s iron hematoxylin stains were performed to visualize elastin fibers as described previously.^58^ For immunostaining, unfixed frozen sections were incubated with acetone for 10 minutes at −20°C. Paraffin-embedded sections (5 µm) were deparaffinized using limonene (#183164, MilliporeSigma). Sections were incubated subsequently with goat serum for 1 hour at 40°C. Rabbit anti-LRP1 (0.5 µg/ml, 15 min, #ab92544, abcam), rabbit anti-α-smooth muscle actin (2 µg/ml, 30 min, #ab5694, abcam), or rabbit anti-PAI1 (for mouse: 0.2 µg/ml, 30 min, #ab222754, abcam; for human: 5 µg/ml, 12 hours, #ab66705, abcam) antibodies were used as primary antibody. Detection of primary antibodies was facilitated with a goat anti rabbit IgG antibody (3 µg/ml, 15 min, BA-1000, Vector) and VECTASTAIN® Elite ABC-HRP Kit (PK-6100, Vector) or ImmPRESS® HRP Goat Anti-Rabbit IgG Polymer Detection Kit (30 min, MP-7451, Vector). ImmPACT AEC (ZH0406, Vector) was used as a chromogen. Isotype matched non-immune rabbit IgG antibody (0.5, 2, or 5 µg/ml, I8140, MilliporeSigma) was used as negative control. Slides were cover slipped with glycerol gelatin (GG1, MilliporeSigma). Antibodies used are described in **Supplemental Table IV**.

Histological images were captured using either Nikon E600 microscope or ZEISS Axio Scan Z1. αSMA or PAI1 positive areas, collagen deposition, and X-gal stained areas were quantified in 40x images (8 bit) using NIS-Elements AR software. Since elastic fibers have autofluorescence illuminated by the FITC (fluorescein isothiocyanate) channel, FITC images of X-gal staining were used to evaluate medial thickening. Medial and adventitial areas were traced using 40x FITC images. X-gal positive area was then assessed in the green channel after subtraction of white color. Elastin fragmentation was defined as the presence of discernable breaks of elastic lamina and counted in the aorta of three serial Movat’s stained sections (100 µm apart). Measurements were verified by an independent investigator who was blinded to study groups.

### Western Blot Analyses

Ascending aortic tissue was harvested from mice and endothelial cells were removed using a cotton swab. Aortic tissues were homogenized in RIPA buffer (#9803, Cell Signaling Technology) with protease inhibitor (#P8340, MilliporeSigma) using Kimble Kontes disposable pellet pestles (#Z359971, DWK Life Science LLC.). For the verification of SHF-specific LRP1 deletion, after the removal of endothelial cells, aortic tissues were incubated with collagenase type I (#SCR13, MilliporeSigma) at 37°C for 12 minutes. Subsequently, adventitia was removed carefully using forceps. Cell lysis buffer (#9803, Cell Signaling Technology) and the protease inhibitor were used for the homogenization. Protein concentrations were determined using DC assay kits (#5000111, Bio-Rad). Equal amounts of protein samples (5 µg) were resolved by SDS-PAGE (10% wt/vol) and transferred electrophoretically to PVDF membranes (#1704273, Bio-Rad). After blocking, antibodies against the following proteins were used to probe membranes: LRP1 (0.4 µg/ml, ab92544, abcam), PAI1 (0.6 µg/ml, ab222754, abcam), and β-actin (1:3000, A5441, MilliporeSigma). Membranes were incubated with either goat anti-rabbit (0.3 µg/ml, #PI-1000, Vector Laboratories) or goat anti-mouse secondary antibodies (1:3000, #A2554, MilliporeSigma). Immune complexes were visualized by chemiluminescence (#34080, Thermo Scientific) and quantified using a ChemiDoc MP Imaging system (#12003154, Bio-Rad).

### Ultrasonography

The ascending aorta was imaged in vivo using a Vevo 2100 ultrasound system with a MicroScan MS550 transducer (40 MHz, FUJIFILM VisualSonics Inc) as described previously.^59, 60^ Briefly, mice were anesthetized using isoflurane (1.0-2.5% vol/vol) and heart rate was adjusted to 400-550 beats per minute during ultrasonography. Aortic luminal diameter was measured between the inner edge to inner edge of the vessel at the end diastole from three separate heart beats. Aortic measurements were verified by an independent investigator who was blinded to study groups.

### Systolic Blood Pressure Measurements

Systolic blood pressure was measured by a non-invasive tail cuff system (Coda 8, Kent Scientific) as described previously.^61^ Conscious mice were restrained in a holder and put on a heated platform. Blood pressure was measured 20 times at the same time each day for three consecutive days. Data showing <60 or >250 mmHg, standard deviation >30 mmHg, or collected cycles <5 of 20 were excluded.

### Aortic Tissue Proteolysis for Mass Spectrometry Assisted Proteomics

Aortic tissues were harvested after 3 days of either saline or AngII infusion, and minced before submersion in RIPA buffer (#9806, Cell Signaling Technology) supplemented with protease inhibitor cocktail (#P8340, MilliporeSigma). Tissue pieces were placed in a Precellys CK14 homogenizing tube with RIPA buffer and ceramic beads (1.4 mm; Bertin Instruments). Samples were homogenized using a Precellys 24 tissue homogenizer using three 10 second cycles at 5,000 rpm. Debris were removed by centrifugation for 10 minutes at 4°C and protein concentrations of supernatant samples were measured using the Pierce BCA Protein Assay (#23225, Thermo Fisher). Equal amounts of protein (10 µg) for each aortic segment were processed using the PreOmics iST in solution trypsinization kit (#00027, PreOmics) according to the manufacturer’s recommended protocols. The final peptide precipitate was dissolved in sample buffer (40 µl, 5% wt/vol acetonitrile, 0.5% wt/vol formic acid in mass spectrometry grade water).

### Mass Spectrometry (MS)

Peptides were diluted [1/2] and analyzed using the Orbitrap Fusion Lumos Tribrid mass spectrometer (Thermo Fisher Scientific) with an Easy-Spray ion source and Easy-nLC1000 HPLC pump (Thermo Fisher Scientific). Peptides were resolved using a dual column set-up: an Acclaim PepMap RSLC C18 trap column, 75 µm x 20 mm; and an EASY-Spray LC heated (45°C) column, 75 µm x 250 mm (Thermo Fisher Scientific). An aqueous to organic gradient (solvent A, 0.1% formic acid in MS-grade water mixed with solvent B, 0.1% formic acid in MS-grade acetonitrile) was generated with a flow rate of 300 nl/min from 5 to 21% solvent B for 75 minutes, 21 to 30% vol/vol solvent B for 15 minutes, followed by ten minutes of a ‘jigsaw wash’, alternating between 5 and 95% vol/vol solvent B. The instrument was set to 120 K resolution, and the top N precursor ions in a 3 second cycle time (within a scan range of 400-1500 m/z; isolation window, 1.6 m/z; ion trap scan rate, normal) were subjected to collision induced dissociation (collision energy 30%) for peptide sequencing (or MS/MS). Dynamic exclusion was enabled (60 seconds).

### MS/MS Data Analysis

The MS/MS data were queried against the mouse UniProt database (downloaded on August 1, 2014) using the SEQUEST search algorithm, via the Proteome Discoverer (PD) Package (version 2.2, Thermo Fisher Scientific), using a 10 ppm tolerance window in the MS1 search space, and a 0.6 Da fragment tolerance window for CID. N-terminal acetylation and methionine oxidation were set as a variable modification, and carbamidomethylation of cysteine residues was set as a fixed modification. In order to quantify peptide precursors detected in the MS1 but not sequenced from sample to sample, we enabled the ‘Feature Mapper’ node. Chromatographic alignment was done with a maximum retention time (RT) shift of 10 minutes and a mass tolerance of 10 ppm. Feature linking and mapping settings were, RT tolerance minimum of 0 minutes, mass tolerance of 10 ppm and signal-to-noise minimum of five. Precursor peptide abundances were based on their chromatographic intensities and total peptide amount was used for normalization. Peptides assigned to a given protein group, and not present in any other protein group, were considered as unique. Consequently, each protein group is represented by a single master protein (PD Grouping feature). We used unique and razor peptides per protein for quantification.

### Aortic Cell Suspension for Single Cell RNA Sequencing (scRNAseq)

Ascending aortic samples were harvested from Mef2c-*Cre* ROSA26R*^mT/mG^* male mice (n=5) at baseline and after 3 days of AngII infusion (1,000 ng/kg/min, H-1705, Bachem, n=4). Aortic samples were pooled in Hanks’ Balanced Salt Solution (HBSS, #14175095, Thermo Fisher Scientific) with fetal bovine serum (10% vol/vol). Periaortic tissues were removed and aortic tissues were cut into small pieces. Aortic samples were subsequently digested with enzyme cocktail (**Supplemental Table V**) in Ca/Mg contained-HBSS (#14025092, Thermo Fisher Scientific) for 60 minutes at 37°C. Cell suspensions were filtered through a 40 μm cell strainer (CLS431750-50EA, MilliporeSigma), centrifuged at 300 g for 10 minutes, and resuspended using cold HBSS (#14175095) with fetal bovine serum (5% vol/vol). Cells were stained with DAPI and sorted to select viable cells (≥ 95% viability) by flow cytometry (FACS Aria III, BD Biosciences). Cells were also sorted based on mTomato and mGFP signals.

### scRNAseq

mGFP positive cells were dispensed onto the Chromium Controller (10x Genomics) and indexed single cell libraries were constructed by a Chromium Single Cell 3’ v3 Reagent Kit (10x Genomics). cDNA libraries were sequenced in a pair-end fashion on an Illumina NovaSeq 6000. Raw FASTQ data were aligned to the Genome Reference Consortium Mouse Build 38 (GRCm38/mm10) reference and gene expressions were quantified using Cell Ranger 3.0 (baseline) or 5.0.1 (AngII).

### Experimental Design

*Randomization:* Each experimental mouse had a unique number generated by the “RAND” function in Excel, and mice were divided into the study groups in numerical order of the unique number.

*Blinding:* All experimental data were verified by an independent investigator blinded to the study group information. For proteomic analyses, the identities of all samples were blinded to the operator.

*Number of replicates:* All experiments included biological replicates. The number of samples in each experiment is described in each figure legend.

*Others:* For all experiments, control data were acquired concurrently with data in which statistical comparisons were performed.

---

**Supplemental Table I.**
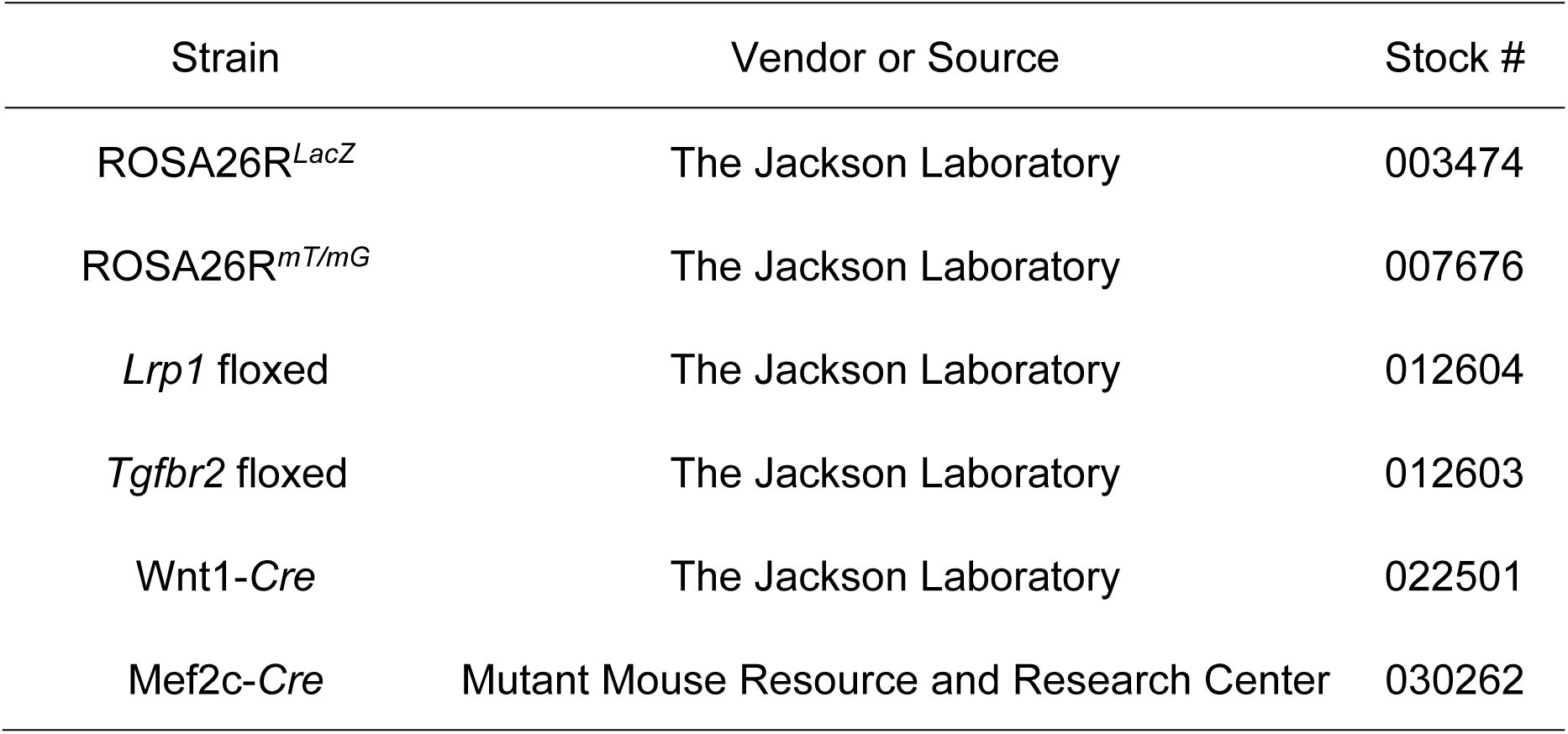
Mouse strains

**Supplemental Table II.** Mouse housing conditions

|  | Mouse Housing Condition | Note |
| --- | --- | --- |
| Set temperature range | 68-74°F (20-23°C) |  |
| Set humidity | 50% |  |
| Light cycle (light:dark) | 14:10 hours |  |
| Water | Reverse osmosis water | ad libitum |
| Feed | 18% Protein Rodent Diet, #2918,<br>Teklad Irradiated Global (Envigo) | ad libitum |
| Bedding | Aspen hardwood chips, #7090A,<br>Harlan Teklad Global (Envigo) |  |
| SPF (Specific pathogen free) | Yes |  |

**Supplemental Table III.**
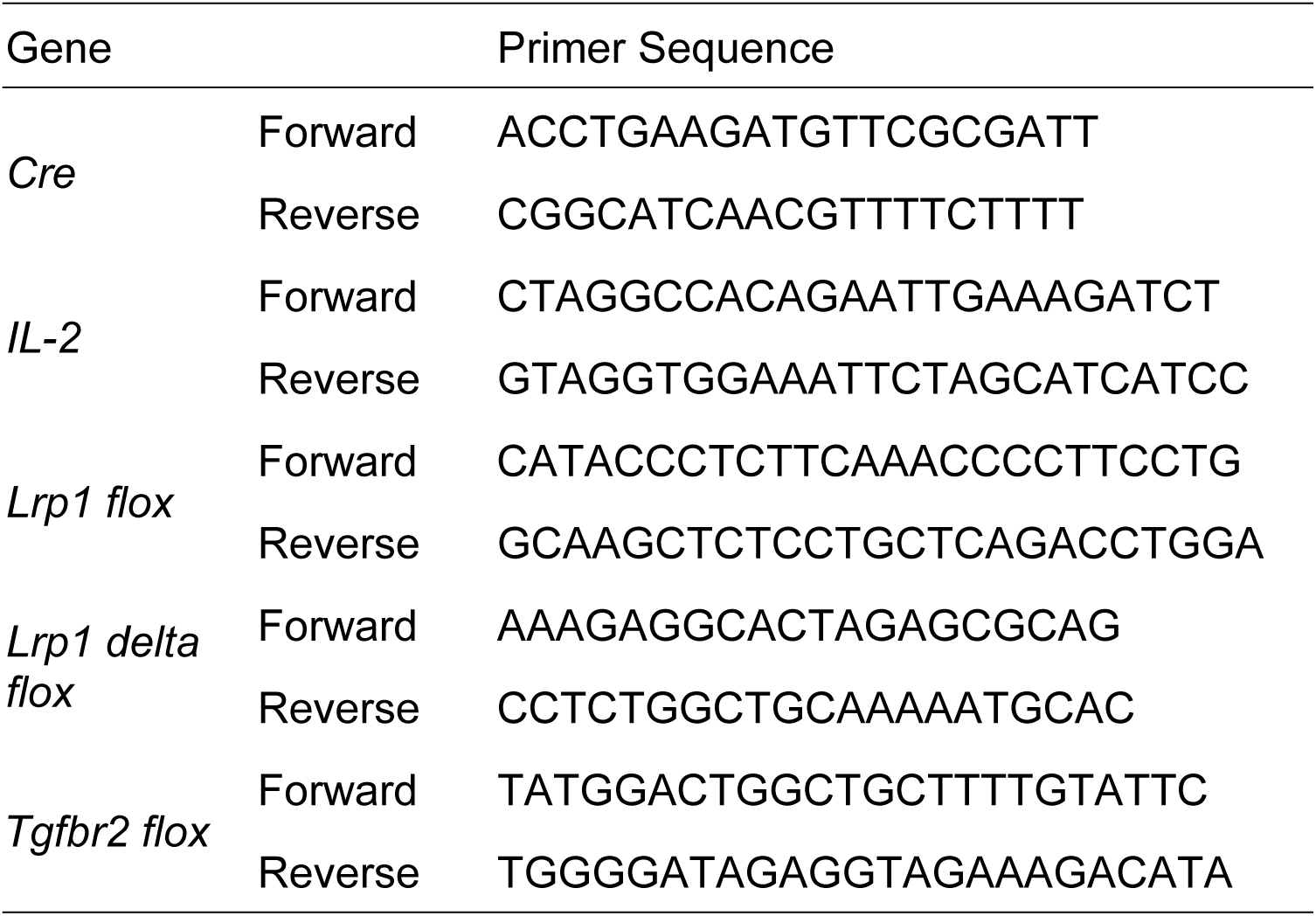
Primer sequences for genotyping

**Supplemental Table IV.** Antibodies

| Target antigen | Vendor | Catalog # | Working Concentration |
| --- | --- | --- | --- |
| Rabbit anti LRP1 | abcam | ab92544 | 0.4 µg/ml for Western blot<br>0.5 µg/ml for immunostaining |
| Rabbit anti α-SMA | abcam | ab5694 | 2 µg/ml |
| Rabbit anti PAI1 | abcam | ab222754 | 0.6 µg/ml for Western blot<br>0.2 µg/ml for immunostaining in mouse aortas |
| Rabbit anti PAI1 | abcam | ab66705 | 5 µg/ml for immunostaining in human TAA tissues |
| Mouse anti mouse β-actin | MilliporeSigma | A5441 | 1:3000 |
| Goat anti-rabbit IgG | Vector Laboratories | PI-1000 | 3 µg/ml |
|  |  | MP-7451 | Ready to use |
| Goat anti-mouse IgG | MilliporeSigma | A2554 | 1:3000 |
| Non-immune rabbit IgG | MilliporeSigma | I8140 | 0.5 µg/ml for LRP1<br>2 µg/ml for α-SMA<br>5 µg/ml for PAI1 |

**Supplemental Table V.**
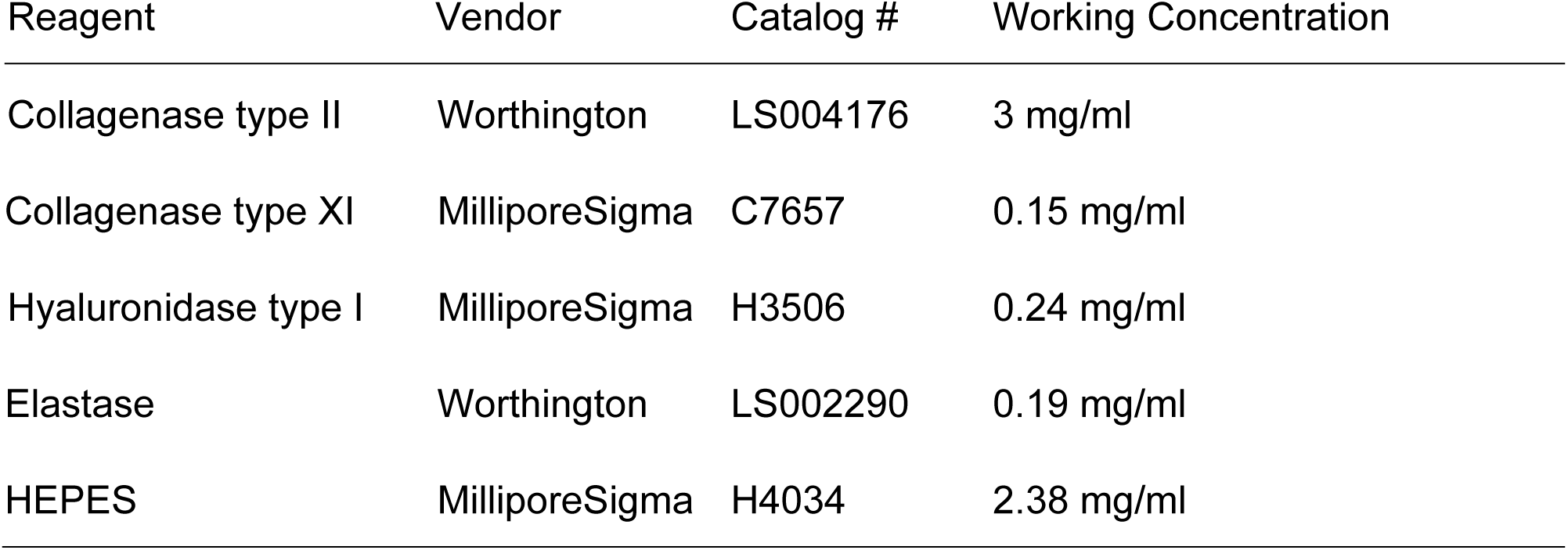
Enzyme cocktail for single cell suspension

| Reagent | Vendor | Catalog # | Working Concentration |
| --- | --- | --- | --- |
| Collagenase type II | Worthington | LS004176 | 3 mg/ml |
| Collagenase type XI | MilliporeSigma | C7657 | 0.15 mg/ml |
| Hyaluronidase type I | MilliporeSigma | H3506 | 0.24 mg/ml |
| Elastase | Worthington | LS002290 | 0.19 mg/ml |
| HEPES | MilliporeSigma | H4034 | 2.38 mg/ml |

**Supplemental Figure I.**
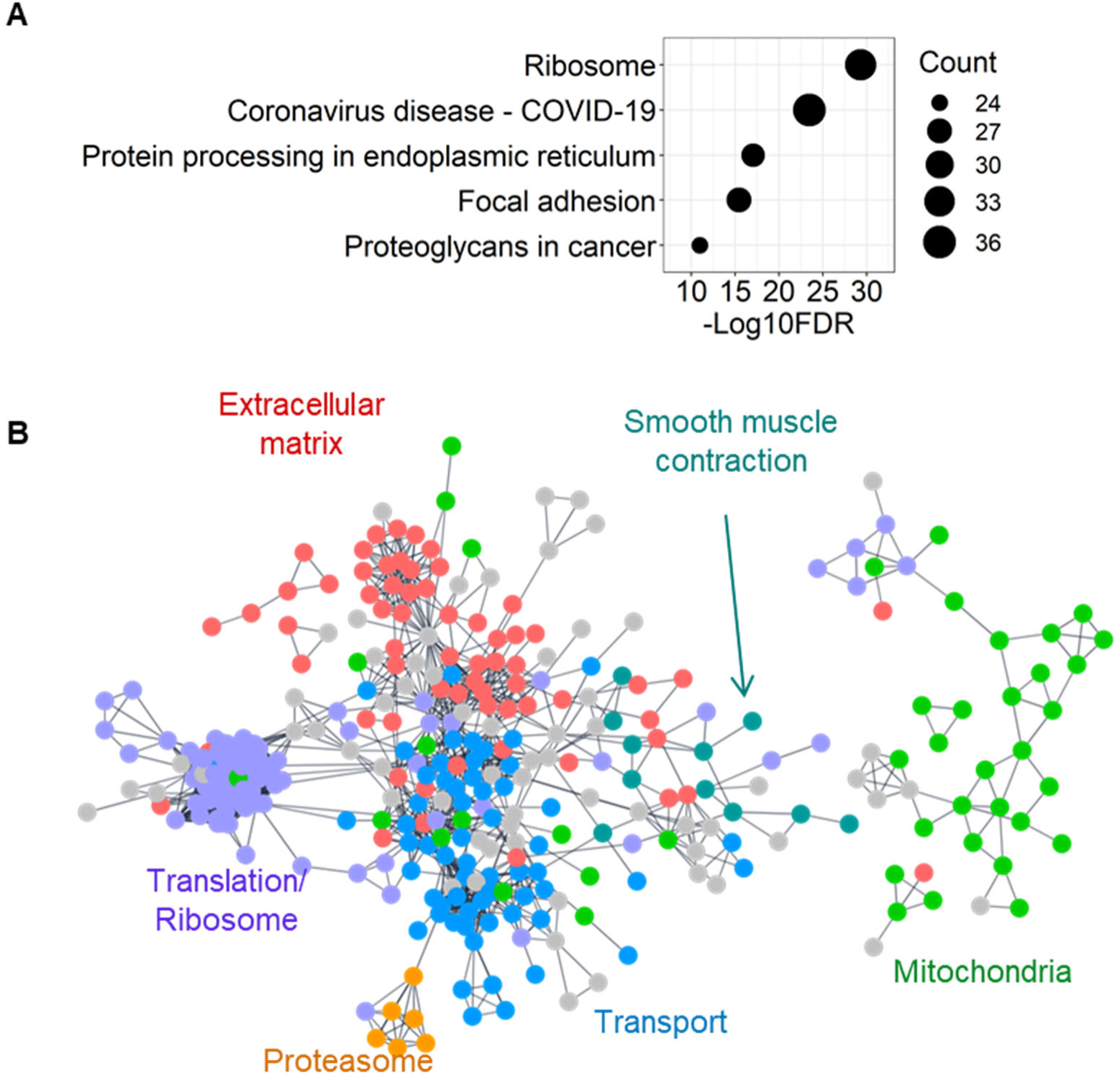
Characteristics of DEGs in the ascending aorta of AngII-infused mice. **(A)** Top 5 significant annotations in enrichment analysis for KEGG pathway of altered molecules detected by the proteomics analysis. **(B)** Protein-protein interaction of 596 altered proteins. Disconnected nodes are not shown.

**Supplemental Figure II.**
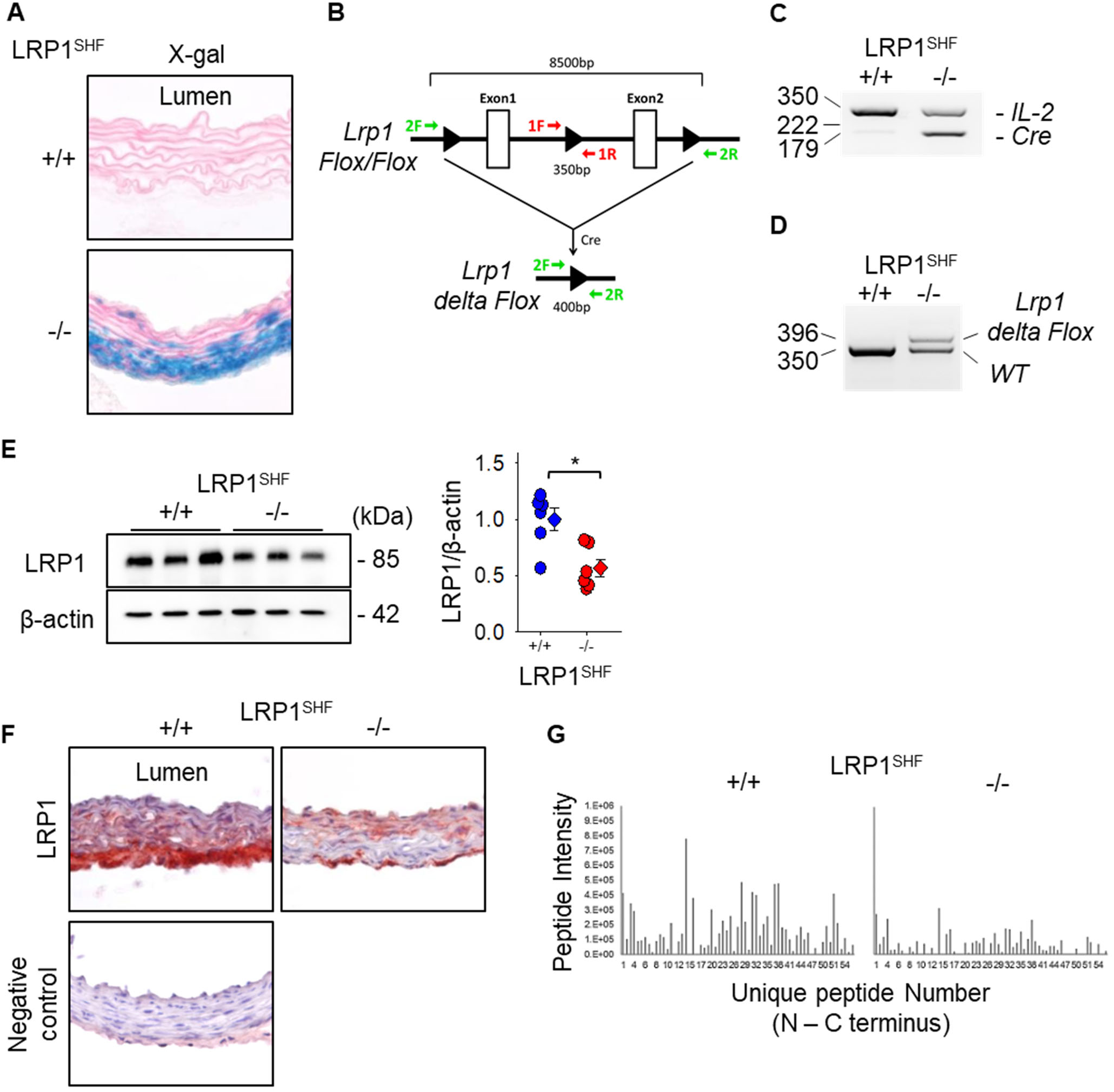
Verification of LRP1 deletion in SHF-derived cells in Mef2c-Cre +/0 *Lrp1* floxed mice. Aortic tissues were harvested from SHF-specific LRP1-deleted mice and wild type littermates at 10 to 12 weeks of age (male, n=3 to 6 per group). **(A)** Representative aortic images of X-gal staining. **(B)** Primer design for PCR of *Lrp1 Flox* sequence. Red and green primers were used for PCR. Representative PCR results of **(C)** *Cre* and **(D)** *Lrp1 delta Flox* sequences. **(E)** Western blot for LRP1 and β-actin and **(F)** immunostaining for LRP1 in the ascending aorta of wild type or SHF-specific LRP1 deleted mice. As negative control, sections were incubated with non-immune rabbit IgG antibody. *P=0.006 by two-tailed Student’s t-test. **(G)** Normalized peptide intensities of LRP1 were exported from Proteome Discoverer 2.2. Peptide intensities were plotted from N to C-terminus with the numbering conserved in the ascending aorta of SHF-specific LRP1-deleted mice and wild type littermates (n=3 per group).

**Supplemental Figure III.**
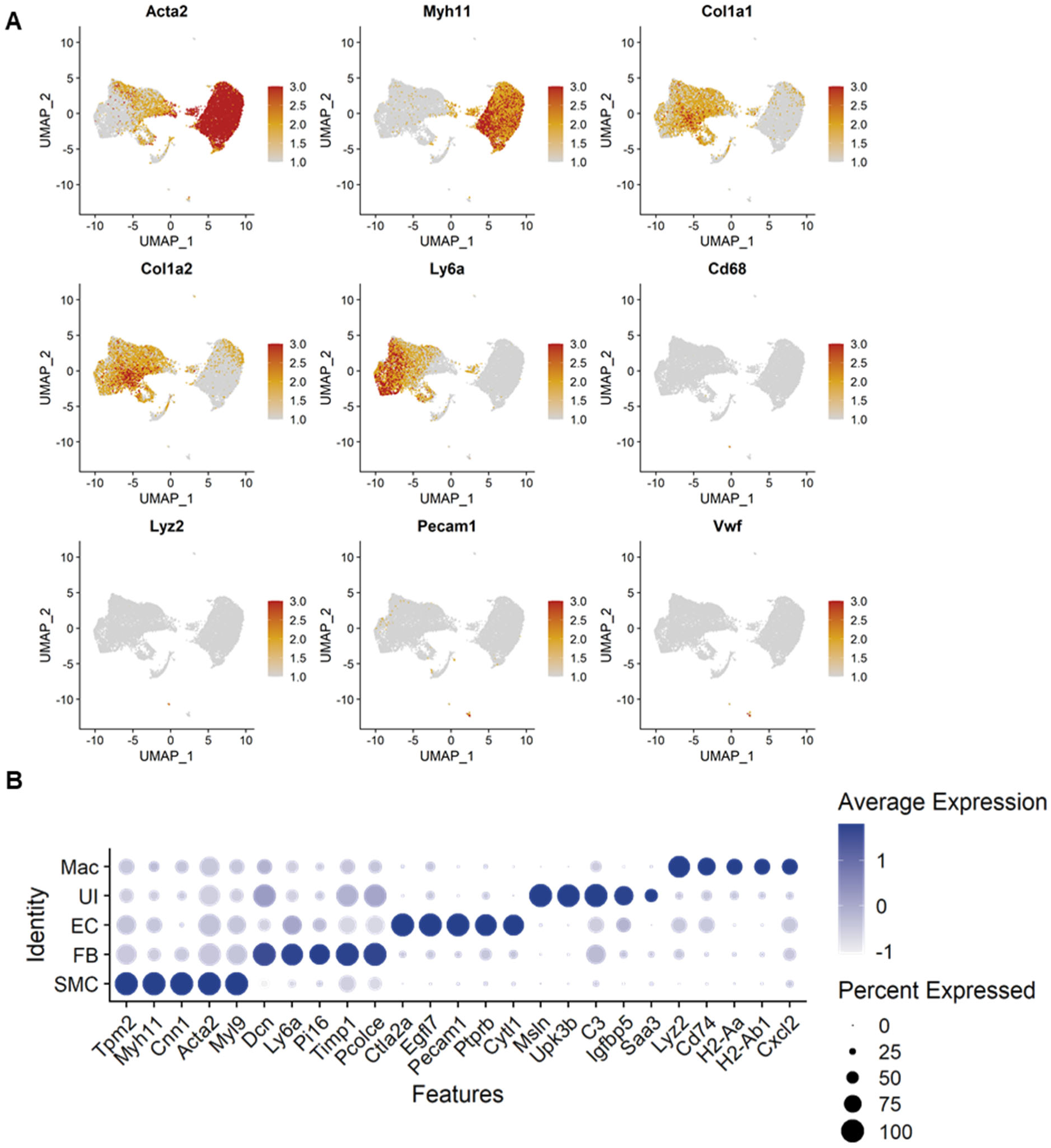
scRNAseq identified multiple cell type clusters in mouse aortas at baseline and after 3 days of AngII infusion. **(A)** Featured plots for cell marker genes in the UMAP plot. **(B)** Dot plots for highly abundance genes in each cell type cluster. SMC indicates smooth muscle cell; FB, fibroblast; EC, endothelial cell; UI, unidentified cell; Mac, macrophage.

**Supplemental Figure IV.**
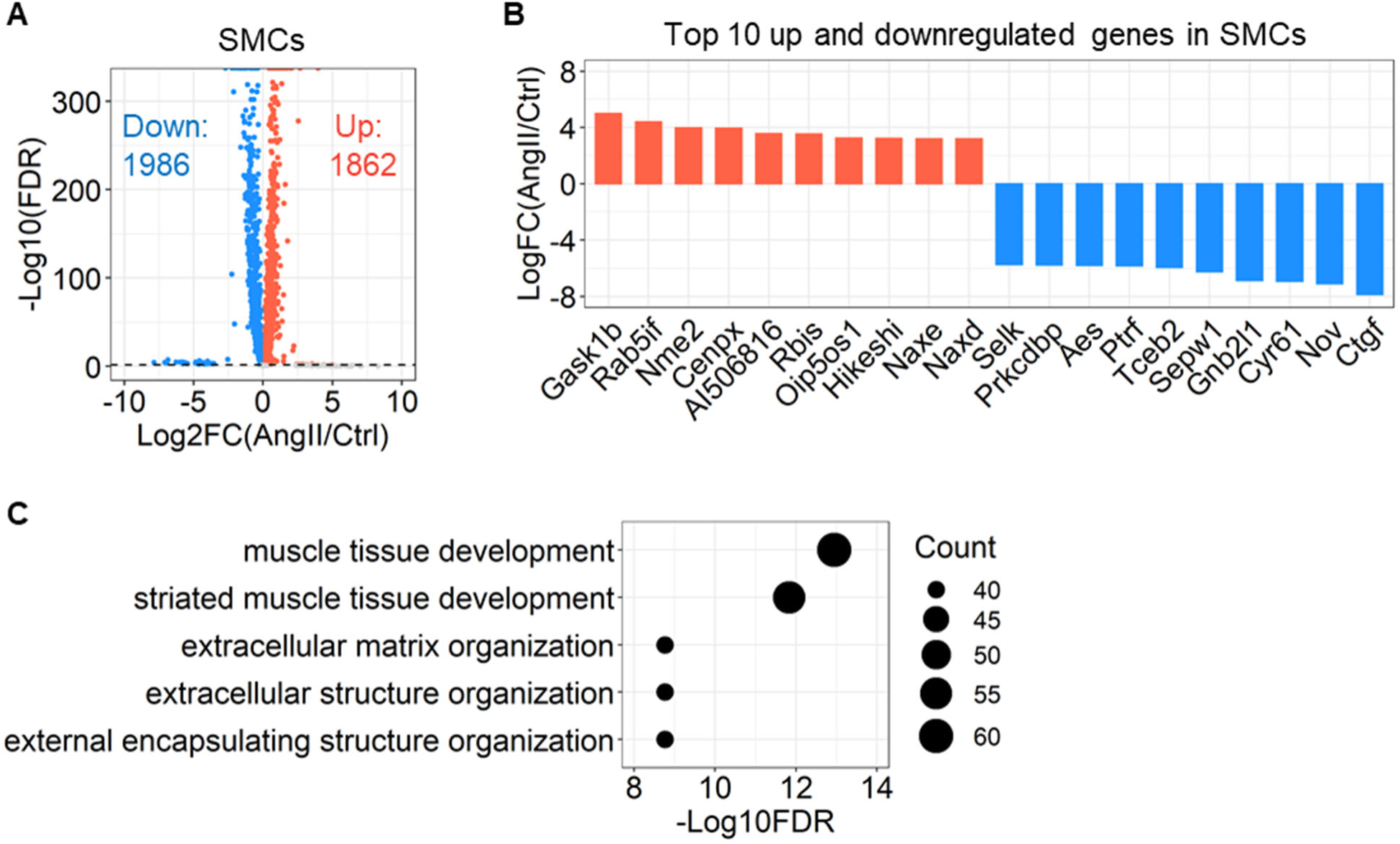
Transcriptomic alteration by AngII infusion in SHF-derived SMCs. **(A)** Volcano plot of DEG analysis, **(B)** top 10 altered genes by AngII infusion, and **(C)** top 5 terms of gene ontology enrichment analysis in the SMC cluster.

**Supplemental Figure V.**
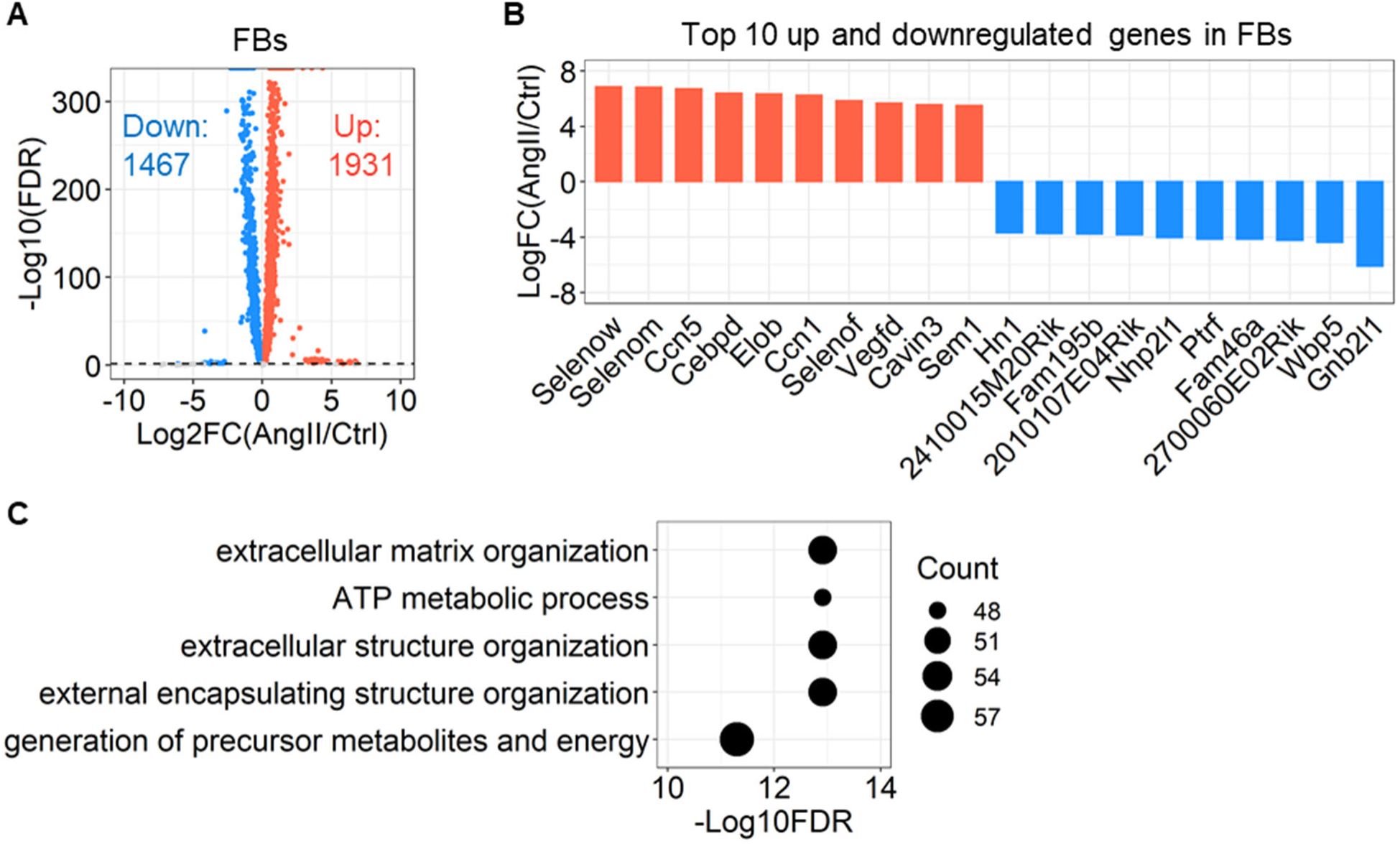
Transcriptomic alteration by AngII infusion in SHF-derived FBs. **(A)** Volcano plot of DEG analysis, **(B)** top 10 altered genes by AngII infusion, and **(C)** top 5 terms of gene ontology enrichment analysis in the FB cluster.

**Supplemental Figure VI.**
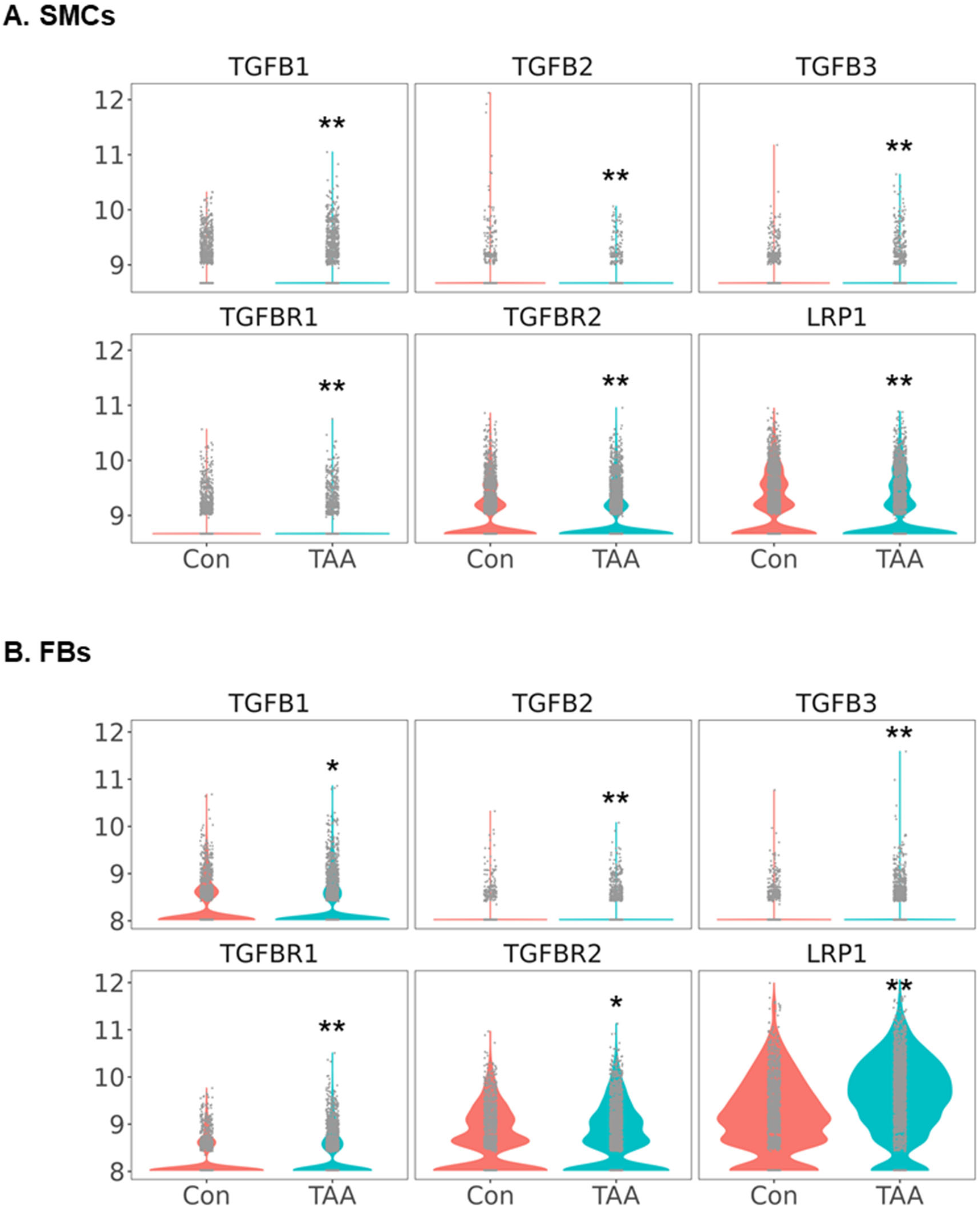
Transcriptomic alteration of TGFβ ligands and receptors in SMCs and FBs of TAAs. Violin plots for TGFβ ligands and receptors in **(A)** SMCs and **(B)** FBs of control and aneurysmal ascending aortas. scRNAseq data were obtained from the GEO (GSE155468). *P<0.001, **P<0.0001 analyzed using the Hurdle model adjested for age implemented.

**Supplemental Figure VII.**
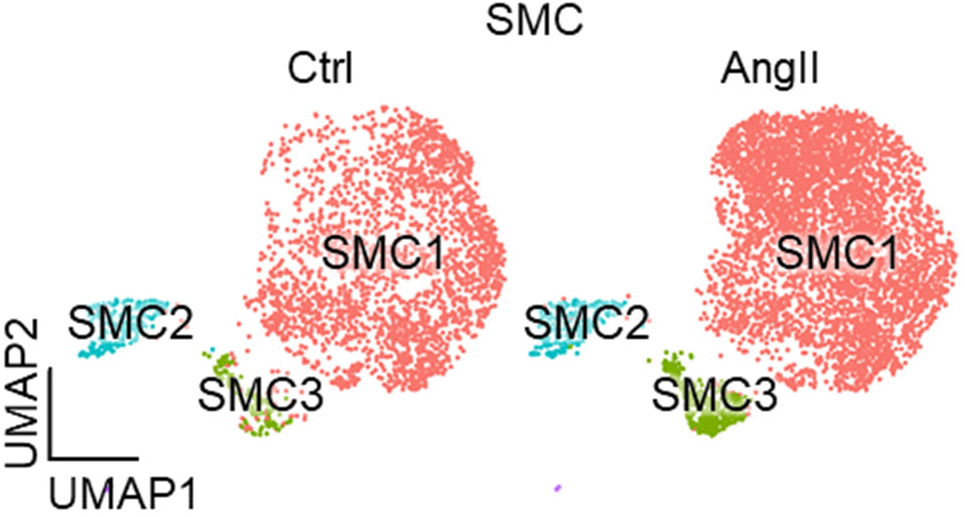
Infusion of AngII did not alter transcriptomic distributions of SMC sub-clusters. UMAP plots at baseline (Ctrl) and after 3 days of AngII infusion in SHF-derived SMCs.

**Supplemental Figure VIII.**
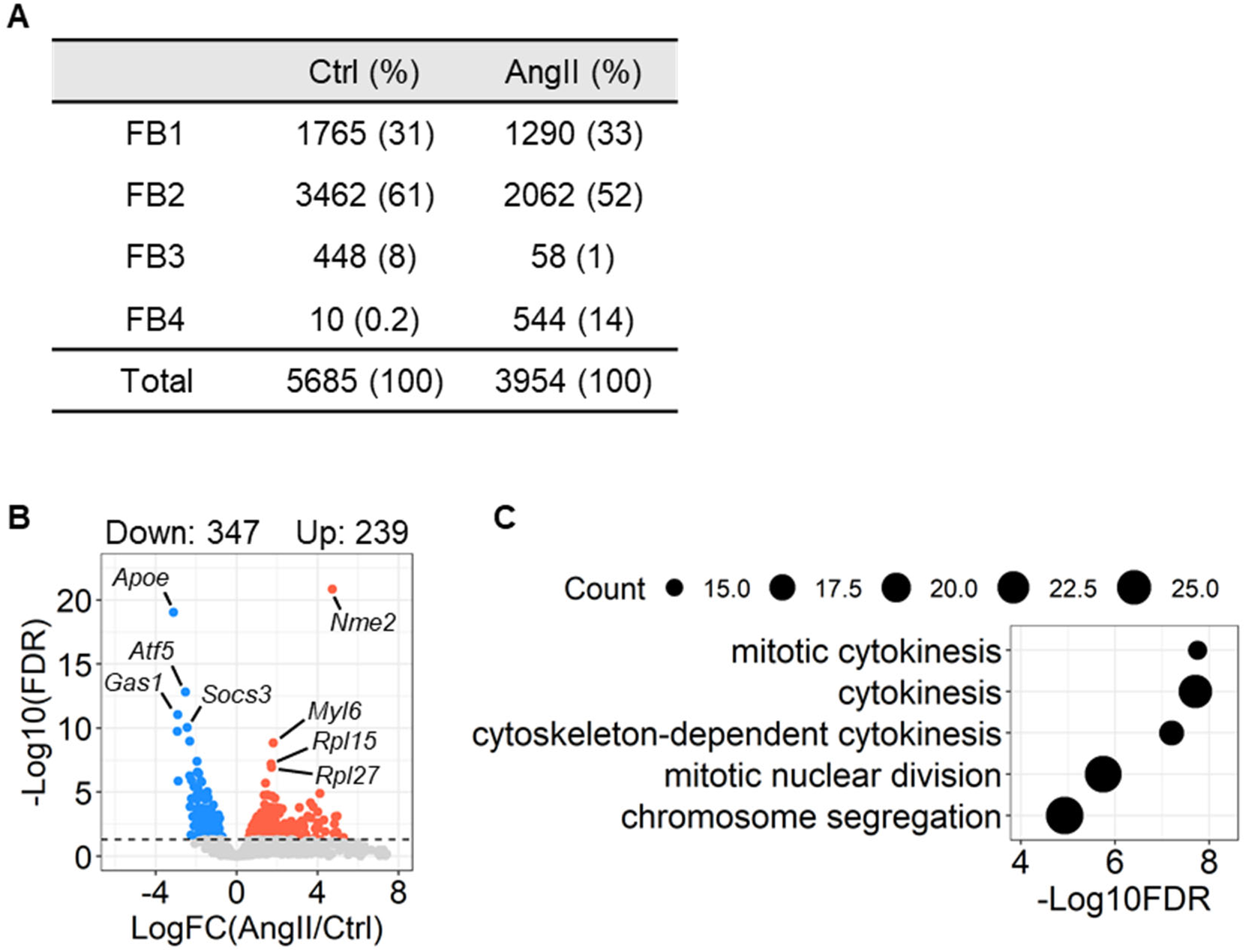
Impact of AngII infusion on the transcriptome of FB4 sub-cluster. **(A)** Number of cells in each FB sub-cluster. **(B)** A volcano plot for DEGs between Ctrl and AngII in FB4 sub-cluster. **(C)** Top 5 significant annotations in enrichment analysis for gene ontology (biological process) using the DEGs in comparing between Ctrl and AngII in FB4 sub-cluster.

**Supplemental Figure IX.**
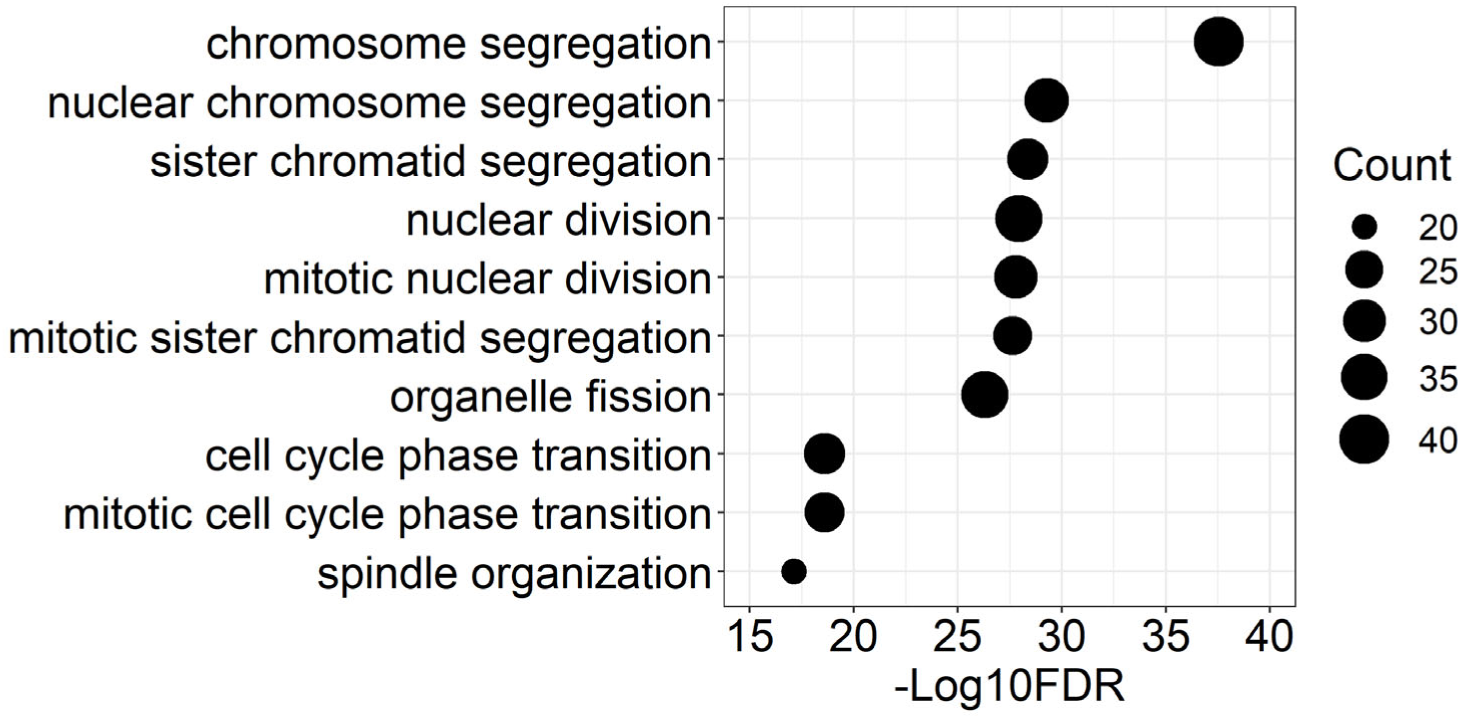
The feature of FB4 sub-cluster. Enrichment analysis for biological process using feature genes expressing in fibroblasts of FB4 sub-cluster.

